# Genomic features of parthenogenetic animals

**DOI:** 10.1101/497495

**Authors:** Kamil S. Jaron, Jens Bast, Reuben W. Nowell, T. Rhyker Ranallo-Benavidez, Marc Robinson-Rechavi, Tanja Schwander

## Abstract

Evolution without sex is predicted to impact genomes in numerous ways. Case studies of individual parthenogenetic animals have reported peculiar genomic features which were suggested to be caused by their mode of reproduction, including high heterozygosity, a high abundance of horizontally acquired genes, a low transposable element load, or the presence of palindromes. We systematically characterized these genomic features in published genomes of 26 parthenogenetic animals representing at least 18 independent transitions to asexuality. Surprisingly, not a single feature was systematically replicated across a majority of these transitions, suggesting that previously reported patterns were lineage specific rather than illustrating general consequences of parthenogenesis. We found that only parthenogens of hybrid origin were characterized by high heterozygosity levels. Parthenogens that were not of hybrid origin appeared to be largely homozygous, independently of the cellular mechanism underlying parthenogenesis. Overall, despite the importance of recombination rate variation for the evolution of sexual animal genomes, the genome-wide absence of recombination does not appear to have had the dramatic effects which are expected from classical theoretical models. The reasons for this are probably a combination of lineage-specific patterns, impact of the origin of parthenogenesis, and a survivorship bias of parthenogenetic lineages.

## Introduction

The switch from sexual reproduction to obligate, female-producing parthenogenesis (thelytoky) has occurred repeatedly among animals and is phylogenetically widespread, with several thousand parthenogenetic animal species described (Bell,1982; van der Kooi et al 2017; Liegeois et al 2020). Parthenogenesis is predicted to have many consequences for genome evolution, since gamete production via meiosis is heavily modified and the restoration of somatic ploidy levels via fertilization no longer takes place. Predicted consequences of parthenogenesis include the accumulation of deleterious mutations [1–3], as well as changes in intragenomic heterozygosity levels [4,5] and transposable element (TE) dynamics [6]. In the present study, we evaluate whether parthenogenesis indeed generates these predicted genomic signatures by reanalyzing and comparing the published genomes of 26 parthenogenetic animal species (**Figure 1**). Previous genome studies were unable to address the question of how parthenogenesis affects genome evolution because they focused on individual lineages. Yet because parthenogenesis is a lineage-level trait, disentangling causes of parthenogenesis from lineage-level characteristics requires replication across independently evolved instances of parthenogenesis. Our study includes species from at least 18 independently evolved parthenogenetic lineages from five different animal phyla, providing us with the unique opportunity to detect general consequences of parthenogenesis that are not solely consequences of lineage-specific evolution. Furthermore, we study the same features in all genomes, whereas many of the original studies focused on different genomic features (**Figure 1**), which thus far precluded broad comparisons across different parthenogenetic groups.

**Figure 1:**
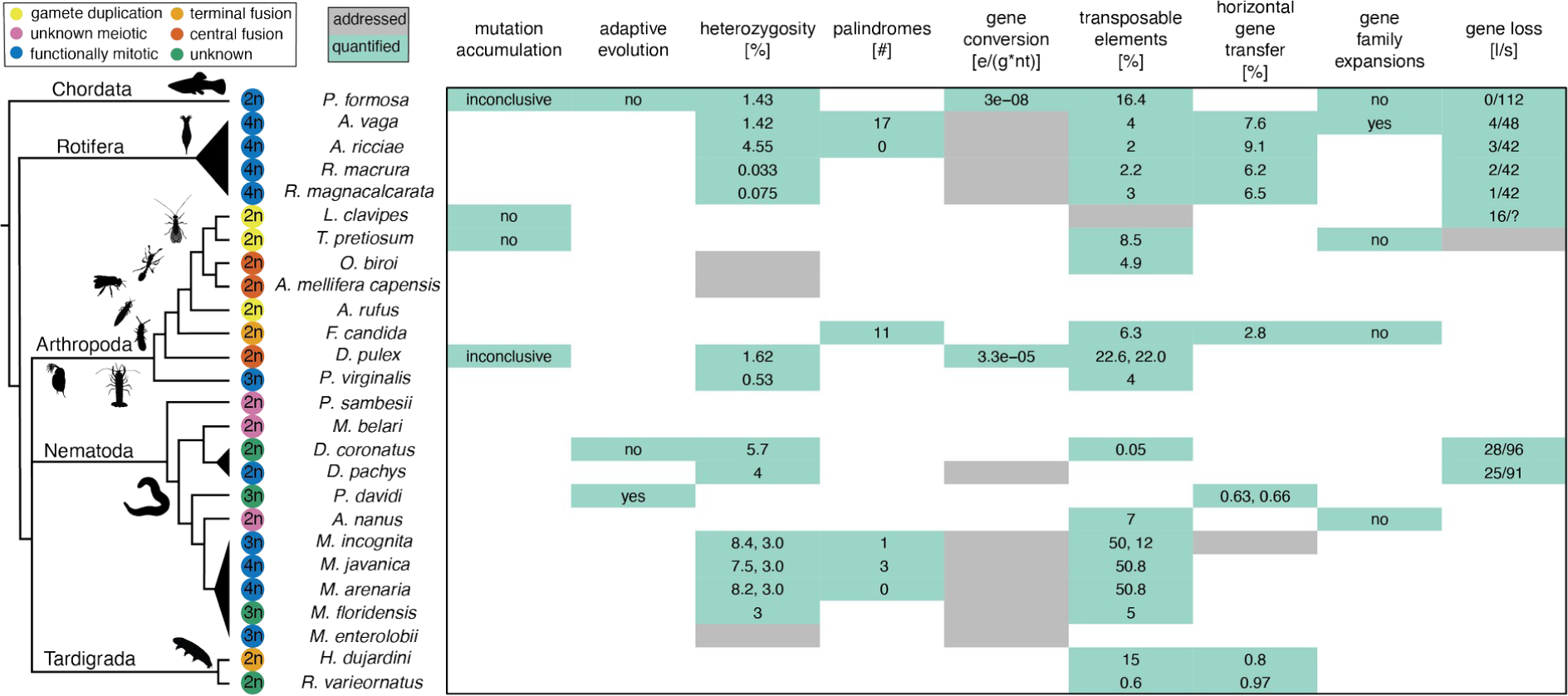
Genome features studied in parthenogenetic animal species. The phylogeny displays the taxonomic relationships of the 26 sequenced parthenogenetic animal species considered here, representing at least 18 independent transitions to parthenogenesis from five different animal phyla. Species that might derive from the same original transition are grouped in triangles. The color of the circle indicates the cellular mechanism of parthenogenesis and the number inside the circle the ploidy of the species (see **Supplemental Table 1** for details). We note *M. floridensis* as triploid, as shown by our analyses, even though it is reported as diploid in the original paper; see **Supplementary Materials S1** for details [11]. Each original genome paper explored a given set of genome features: the green cells represent cases where the genomic feature was quantified (values are indicated); the grey cells represent studies where the genomic features were addressed with respect to parthenogenesis, but the results were not quantitatively comparable to other studies. We reanalysed heterozygosity, palindromes, transposable elements, and horizontal gene transfer in this study; the discussion of the remaining features is based on the analyses reported in the individual genome studies [12–35]. Findings for mutation accumulation and adaptive evolution refer to comparisons between sexual and asexual species and are reported with respect to theoretical predictions (yes: as predicted, no: opposite to predictions, inconclusive: no difference). e/(g*nt): **e**vent per **g**eneration per **n**ucleo**t**ide; l/s: number of **l**ost genes among the **s**tudied genes related to sexual reproduction.

Because the predicted consequences of parthenogenesis are strongly affected by how asexuality evolved from the sexual ancestor (**Box 1**) as well as by the cellular mechanisms underlying parthenogenesis (**Box 2**), we include biological differences among species in our comparisons. For example, some parthenogenetic species have evolved via hybridization (**Box 1**), which generates incipient parthenogens with high intragenomic heterozygosity and can result in increased activity of transposable elements [7–9]. In such instances, it can be difficult to disentangle consequences of hybridization from those of parthenogenesis. Similarly, some cellular mechanisms underlying parthenogenesis involve meiotic divisions, with a secondary restoration of somatic ploidy levels, while others do not. In the former case, heterozygosity is expected to decay rapidly, while in the latter case, it could be maintained or even increase over time [10]. Finally, because the genome studies differed in their focus and in the methods used, we reanalyzed the published genomes with standardized approaches.

Whenever possible, we conducted quantitative comparisons between groups of parthenogenetic species. However, for interpretation, it is important to consider that the available genomes are neither a random nor a fully representative sample of parthenogenetic animals.

Using the 26 genomes, we studied nine genomic features that have been proposed to be affected by parthenogenesis. Four of them represent classical theoretical predictions for consequences of asexuality on genome evolution, namely, consequences for mutation accumulation, positive selection, transposable element dynamics, and intragenomic heterozygosity (see below). The five remaining ones are unusual genomic features that were observed in individual parthenogenetic species and suggested to be linked to their mode of reproduction: horizontally acquired genes, palindromes, gene conversion, gene family expansions, and gene losses, see below.

We first reviewed the literature for information on all nine genomic features in the 26 parthenogenetic species (**Figure 1**). However, the methods used to evaluate a given genomic feature varied among studies of parthenogenetic species (**Supplementary Figure 3**), which precluded quantitative comparisons. Hence we developed a standardized pipeline to quantify heterozygosity, transposable element load, the frequency of horizontally acquired genes, and palindromes. Heterozygosity and transposable element loads were quantified using raw sequencing reads to avoid biases introduced by differences in genome assembly quality, while analyses of horizontally acquired genes and palindromes were based on the published genome assemblies because of methodological requirements. We relied on published information (without reanalysis) for evaluating mutation accumulation, positive selection, gene family expansions, gene losses and gene conversion because these genome features cannot be studied using individual genomes of parthenogenetic species. Indeed, analysing these features requires genomes of sexual relatives and/or population genomic data which is not available for the vast majority of the 26 parthenogenetic species.

### Box 1

Transitions to parthenogenesis

Meiotic sex and recombination evolved once in the common ancestor of eukaryotes [36]. Parthenogenetic animals therefore derive from a sexual ancestor, but how transitions from sexual to parthenogenetic reproduction occur can vary and have different expected consequences for the genome [8].

#### Hybrid origin

Hybridization between sexual species can generate hybrid females that reproduce parthenogenetically [8,37]. Parthenogenesis caused by hybridization can generate a highly heterozygous genome, depending on the divergence between the parental sexual species prior to hybridization. Hybridization can also result in a burst of transposable element activity [7].

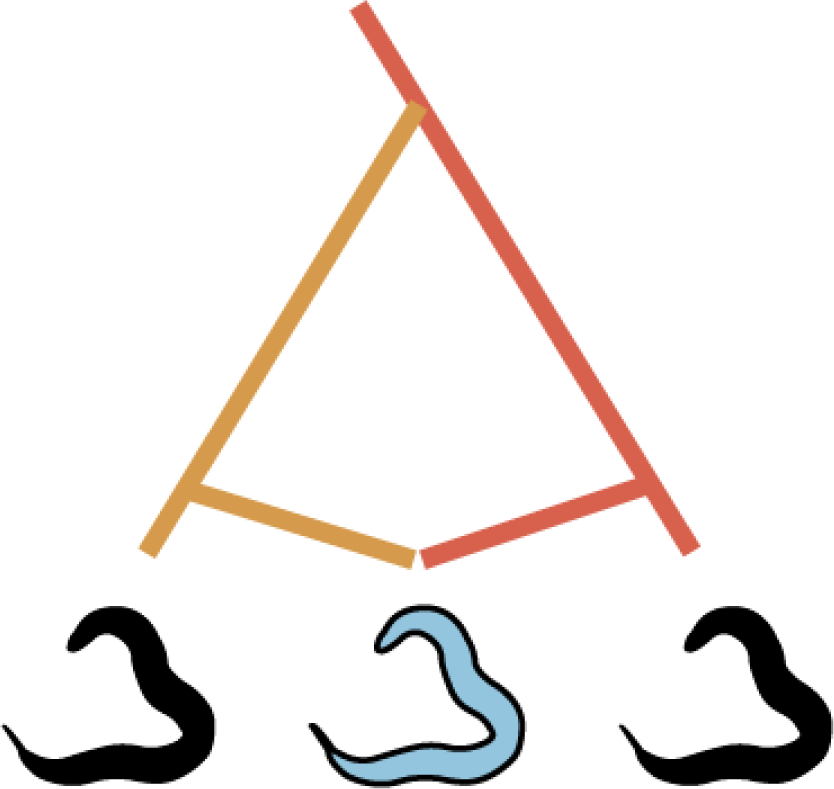

#### Intraspecific origins

##### Endosymbiont infection

Infection with intracellular endosymbionts (such as *Wolbachia, Cardinium* or *Rickettsia*) can cause parthenogenesis, a pattern that is frequent in species with haplodiploid sex determination [38]. This type of transition often (but not always) results in fully homozygous lineages because induction of parthenogenesis frequently occurs via gamete duplication (see **Box 2**).

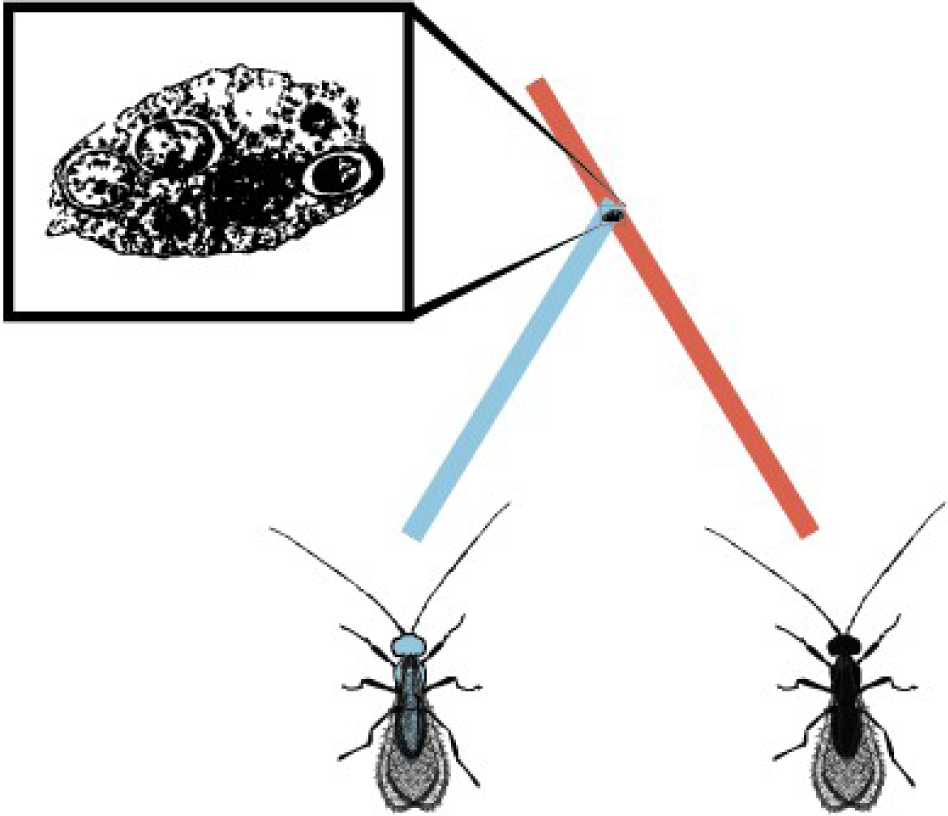

##### Spontaneous mutations/Contagious parthenogenesis

Spontaneous mutations can also underlie transitions from sexual to parthenogenetic reproduction. In addition, parthenogenetic females of some species produce males that mate with females of sexual lineages, and thereby generate new parthenogenetic strains (contagious parthenogenesis). In both cases, the genomes of incipient parthenogenetic lineages are expected to be very similar to those of their sexual relatives and subsequent changes should be largely driven by the cellular mechanism underlying parthenogenesis (**Box 2**).

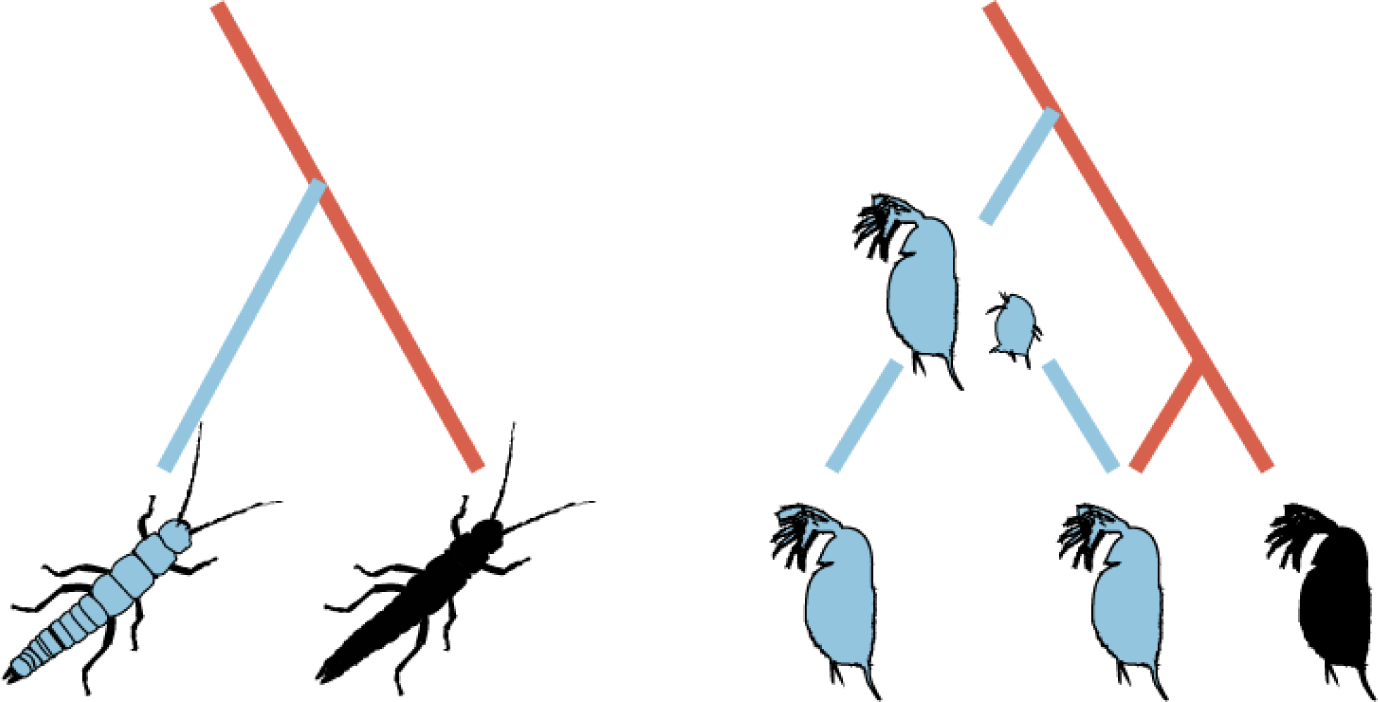

### Box 2

Cellular mechanisms of parthenogenesis

In sexual species offspring are generated through the fusion of male and female gametes. In parthenogens, females generate diploid (or polyploid) offspring from unfertilized oocytes via different cellular mechanisms. The mechanism is predicted to affect genome evolution and especially heterozygosity levels. For details see [10,39].

#### Functionally mitotic parthenogenesis (Apomixis)

Under functionally mitotic asexuality, no ploidy reduction occurs and offspring are clones of their mother. Note that because gametogenesis and egg production are tightly linked to meiotic divisions in sexual species, functionally mitotic parthenogenesis derives from meiosis and does therefore generally not correspond to mitotic divisions in terms of molecular mechanisms. Many functionally mitotic parthenogens notably still feature vestiges of the ancestral meiotic divisions such as homologous chromosome pairing or chromosome condensation typical for meiosis.

#### Meiotic parthenogenesis (Automixis)

Under meiotic parthenogenesis, meiotic divisions occur partially or completely, but somatic ploidy levels are maintained via different mechanisms. Depending on how recombination occurs, some of these mechanisms are equivalent to functionally mitotic parthenogenesis, even though meiosis is fully maintained.

##### Endoduplication

A duplication of the entire chromosome set occurs before normal meiosis, during which ploidy is reduced again. If recombination occurs between identical chromosome copies rather than between chromosome homologs, endoduplication is functionally mitotic, i.e., produces offspring that are clones of their mother.

##### *Central fusion* and *terminal fusion*

Under these two mechanisms, somatic ploidy levels are restored through the fusion of two of the four meiotic products (products separated during the first meiotic division merge under central fusion, products separated during the second division merge under terminal fusion). In the absence of recombination, central fusion generates offspring that are clones of their mother, while terminal fusion generates fully homozygous offspring. The consequences for heterozygosity are opposite under inverted meiosis, where chromatids are separated during meiosis I and chromosomes during meiosis II. For example, terminal fusion with an inverted sequence of meiosis and no recombination (not shown here) generates offspring that are clones of their mother (see [40] for a recent review).

##### Gamete duplication

After a full meiosis, a haploid meiotic product undergoes duplication. This results in a diploid, but fully homozygous offspring.

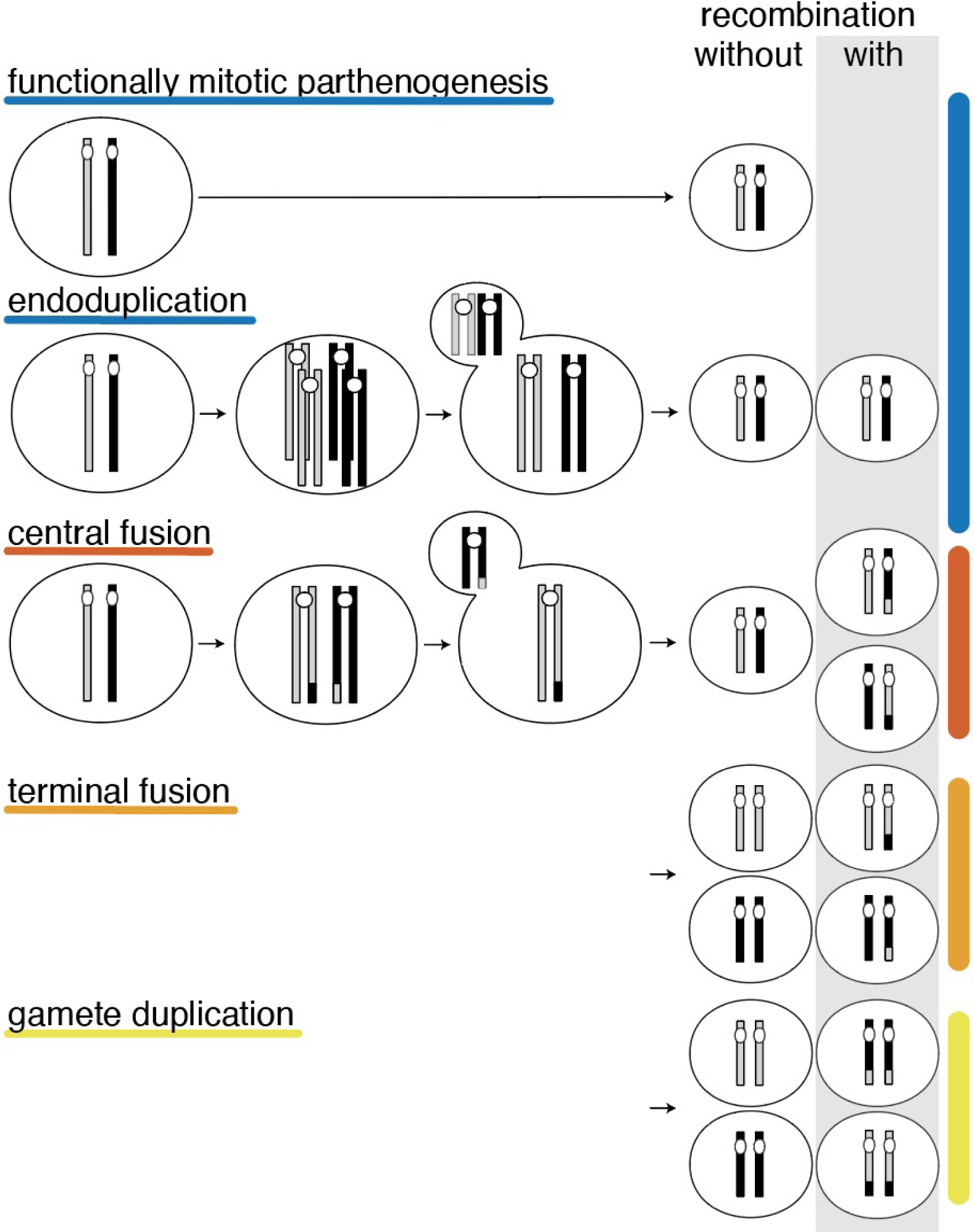

## Methods

### Overview of species and genomes studied

We reanalyzed the published genomes of 26 asexual animal species with the aim of identifying general genomic signatures of asexualilty. The 26 species correspond to at least 18 independent transitions to asexuality and cover a broad taxonomic range, including chordates, rotifers, arthropods, nematodes, and tardigrades. In addition to covering this taxonomic range, these asexual species vary in the cellular mechanisms underlying parthenogenesis, in the mechanisms that caused the transition to asexuality, ploidy, as well as in other biological aspects (**Figure 1, Supplementary Tables 1 & 2**). This variation allows us to assess whether parthenogenesis generates universal genomic signatures independently of species-specific traits.

The cellular mechanisms underlying parthenogenesis have been reported in 22 of the 26 species. Eleven of them reproduce via functionally mitotic parthenogenesis, while the 11 remaining species have different types of meiotic parthenogenesis (**Figure 1**). Information on how asexuality evolved is available for 16 of the 26 sequenced species (**Supplementary Table 1**). A hybrid origin has been suggested for ten of these, based on identification of parental species and/or genomic analyses. Endosymbionts are the most likely cause of asexuality in four species (the springtail, both wasps, and the thrips), and spontaneous mutation in two (the ant and the cape honey bee).

Most if not all predicted consequences of asexuality are expected to accumulate over time, meaning that their effect size as well as the power to detect them is increased in old asexual lineages. However, estimating the age of asexual lineages is difficult and always associated with large uncertainties [42,43]. Therefore we did not include quantitative comparisons among asexuals with respect to their age. However, because our set of species comprises some asexuals believed to be ‘ancient’ (i.e., several million years old, see **Supplementary Table 1**), we discuss, where appropriate, potential age effects in a qualitative manner.

### Recomputed genomic features

We combined different methods into a complete pipeline that collects published assemblies, sequencing reads, and genome annotation data from online databases, and automatically computes the focal genome features. The pipeline is available at https://github.com/KamilSJaron/genomic-features-of-parthenogenetic-animals. We provide a summary of each method below, with technical details (e.g., specific parameters used for each method) indicated in the **Supplementary Methods**. For some species, additional genomes to the ones we used were available, but we did not include them because of low data quality and/or unavailable Illumina reads (this was the case for one sample of *M. incognita, M. floridensis* and multiple samples of *D. pulex* [22,30,44]). The accession numbers of the individual sequencing projects are listed in **Supplementary Table 1**.

### Heterozygosity

To compare intragenomic heterozygosity among species with different ploidy levels, we estimated heterozygosity as the proportion of sites with more than one allele present among all homologous genome regions (consistent with [45]). In diploid species, this genome-wide heterozygosity can correspond to the divergence between alleles (homolog heterozygosity), or if the species has a history of hybridization, to the divergence of gene copies derived from different species (hereafter homoeologs, following the terminology of Glover et al [57]). In polyploid species, heterozygosity can be a combination of homolog and homoeolog divergence. We distinguished homolog and homoeolog heterozygosity whenever possible, or inferred a “composite heterozygosity” (the sum of the two) otherwise. To avoid biases stemming from variable genome assembly qualities, we estimated heterozygosity directly from sequencing reads using kmer spectra analysis [11], except for four bdelloid rotifers where the heterozygosity levels between homoeologs exceeded the range quantifiable by this method (see **Supplementary Materials S2**).

Kmer spectra analysis is based on the decomposition of sequencing reads into all possible subsequences (kmers) and generating a coverage histogram of the kmers (kmer spectrum). We performed the kmer spectra analysis using GenomeScope 2.0 [11]. Genomescope fits a mixture model of evenly spaced negative binomial distributions to the kmer spectrum. Fits are then used to estimate heterozygosity. With the exception of bdelloid rotifers, we are not able to directly compare the divergence of individual haplotypes (because such a comparison requires phased genomes). However, we are able to measure the haplotype structure on a per-locus basis. These approaches notably allow us to distinguish biallelic from triallelic loci in triploid organisms.

### Transposable elements

We quantified transposable elements using DnaPipeTE [46]. This method uses the haploid genome size to subsample sequencing reads to a low coverage of 0.5x. These subsampled reads are then assembled using an assembler (Trinity) that can deal with uneven coverages and is able to split assembled regions with few differences (including different TE families). The assembled sequences largely correspond to repetitions as non-repetitive genome regions present in the subsampled reads drop out at this stage, because the coverage of such regions is too low for assembling. The assembled sequences are annotated by homology using a database of known TEs. The output of the method is the number of sampled nucleotides assembled and annotated as different types of repeats, and fractions are calculated as the numbers divided by the total number of sampled nucleotides. Our reported values of TE loads include only repeats that were annotated as TEs, i.e., we did not include ‘unknown’ repeats which consist of tandem repeats (satellite repeats), duplications or very divergent/unknown TEs. Note that in addition to the 26 parthenogenetic genomes (Figure 1), we also analysed one sexual species *Procambarus fallax* (SAMN06115719) for comparison to its asexual sister species *P. virginalis*.

### Palindromes

Palindromes are formed of two homologous reverse complementary sequences on the same chromosome (**Supplementary Figure 2**). Palindromes can facilitate gene conversion and therefore help to escape mutational meltdown via Muller’ s ratchet [47,48]. To test if they play such a role in asexual organisms we identified palindromes using colinearity analysis imlplemented in the program MCScanX [49]. Detected collinear blocks were filtered to contain only reverse complementary collinear blocks on the same chromosome, since only such structures have the capacity to form a hairpin (**Supplementary Figure 2**).

This method for palindrome identification depends on genome assemblies. Palindromes are less likely to be detected in highly fragmented assemblies, and artificial palindromes can be generated by erroneous scaffolding (see also [14]). Our analyses assume that there are no systematic scaffolding errors in the published assemblies, meaning that our list of palindromes includes false positives that are generated by mis-assemblies in the published reference genomes. Palindrome identification methods rely on genome annotations, which are available for 23 of the 26 asexual species (all except *D. pulex, A. mellifera capensis*, and *A. rufus*). We screened these 23 genomes for the presence of palindromic arrangements (See **Supplementary Methods** for details).

### Horizontal Gene Transfer

We systematically estimated the percentage of non-metazoan HGT candidates (HGTC) in the 23 of the 26 asexual species with available gene annotations using a sequence comparison based approach, following [14]. For each species, we compared the set of annotated genes to the UniRef90 and UniProtKB/Swiss-Prot protein databases to identify genes of likely non-metazoan origin [50]. We considered non-metazoan genes as HGT candidates only if they were on a scaffold that also encoded at least one gene of unambiguous metazoan origin, to control for potential contamination in the genome assemblies (see **Supplementary Methods** for details).

## Results and discussion

There has been an accumulation of genome studies of individual parthenogenetic animal species (**Figure 1**) and most of these studies have suggested that one or a few specific genome features uncovered in their focal species might be linked to parthenogenesis. To test whether any of these genomic features were in fact general consequences of parthenogenetic reproduction rather than lineage-specific traits, we evaluated evidence from published genomes of 26 parthenogenetic animals in the light of theoretical expectations for genome evolution under parthenogenesis. We quantified heterozygosity, TE loads, the frequency of horizontally acquired genes and palindromic sequence arrangements using standardized methods, and combined these analyses with a review of published information on mutation accumulation, positive selection, gene family expansions, and gene losses.

### Mutation accumulation and positive selection

One of the classical predictions linked to parthenogenesis is that it reduces the efficacy of selection [1–3,51–53]. This reduction occurs because linkage among loci in asexual species prevents selection from acting individually on each locus, resulting in different forms of selective interference [54]. This selective interference can result in a faster accumulation of deleterious mutations and a slower rate of adaptation. While there is accumulating evidence for these processes in experimental evolution studies (e.g., [55–57]), their impact for natural populations remains unclear [58]. Genomic data can provide the basis for studying mutation accumulation and adaptation (positive selection) in natural populations.

In summary, results from genome-wide studies addressing the prediction of deleterious mutation accumulation in natural populations of parthenogenetic species are equivocal. More studies are therefore needed. A major constraint for studying deleterious mutation accumulation, and the reason why it was not studied in most genome studies of parthenogenetic species (**Figure 1**), is that it requires homologous gene sets from sexual outgroups for comparison. These species are either unknown or not included in most published genome studies of parthenogens.

The same constraints likely explain why no study has thus far directly addressed adaptive evolution in the genome of a parthenogenetic species. The question of adaptive evolution was addressed indirectly in the amazon molly, by studying the amount of segregating variation at immune genes (where variation is known to be beneficial). The authors found very high diversities at immune genes [12]. However, these were difficult to interpret because standing variation was not compared to that in sexual relatives, and because the amazon molly is a hybrid species. Hence the high diversity could be a consequence of the hybrid origin rather than of parthenogenesis. Furthermore, sexual and parthenogenetic sister species differ in other aspects besides their reproductive mode, which complicates interpretations of such individual comparisons.

### Heterozygosity

Intragenomic (individual-level) heterozygosity is the nucleotidic divergence between the haploid genome copies of an individual. In a panmictic sexual population, the level of intragenomic heterozygosity corresponds to the level of genetic diversity in a population (the amount of variation observed between DNA sequences from different individuals). This is however not the case in asexual populations, which are, by definition, not panmictic [5].

Intragenomic heterozygosity in asexual organisms is expected to depend on three major factors: (1) the mechanism of transition to parthenogenesis (which determines the initial level of heterozygosity, expected to be high for hybrid origins; **Box 1**), (2) the cellular mechanism underlying parthenogenesis (which determines whether heterozygosity will increase or decrease over time; **Box 2**), and (3) how long a species has been reproducing asexually (because the effect of parthenogenesis accumulates over time).

As expected, all of the species with a known hybrid origin of parthenogenesis display high heterozygosity levels (1.73% - 8.5%, **Figure 2**). By contrast, species with an intraspecific origin of parthenogenesis show low heterozygosity levels (0.03% - 0.83%, **Figure 2**). In addition to hybrid origins, polyploidy may also contribute to high heterozygosity in parthenogens as heterozygosity is higher in polyploid (1.84% - 33.21%) than diploid species (0.03% - 5.26%). It is impossible to disentangle the effects of hybrid origin from polyploidy on heterozygosity in our dataset as across the 26 species, hybrid origins are correlated with polyploidy. Six of the 11 polyploids in our sample are of hybrid origin, while for the five others a hybrid origin is supported by our results (see below), even though it was not suggested previously. While the correlation between hybrid origin and polyploidy in our dataset is striking, it is important to note that this correlation would most likely be weaker in a random sample of parthenogenetic animals. Indeed, many polyploid parthenogenetic animals are not of hybrid origin, including several well studied species such as the New Zealand mudsnail *Potamopyrgus antipodarum*, the bush cricket *Saga pedo*, or the bagworm moth *Dahlica triquetrella*. None of these has a published genome yet, which precludes their inclusion in our study.

**Figure 2:**
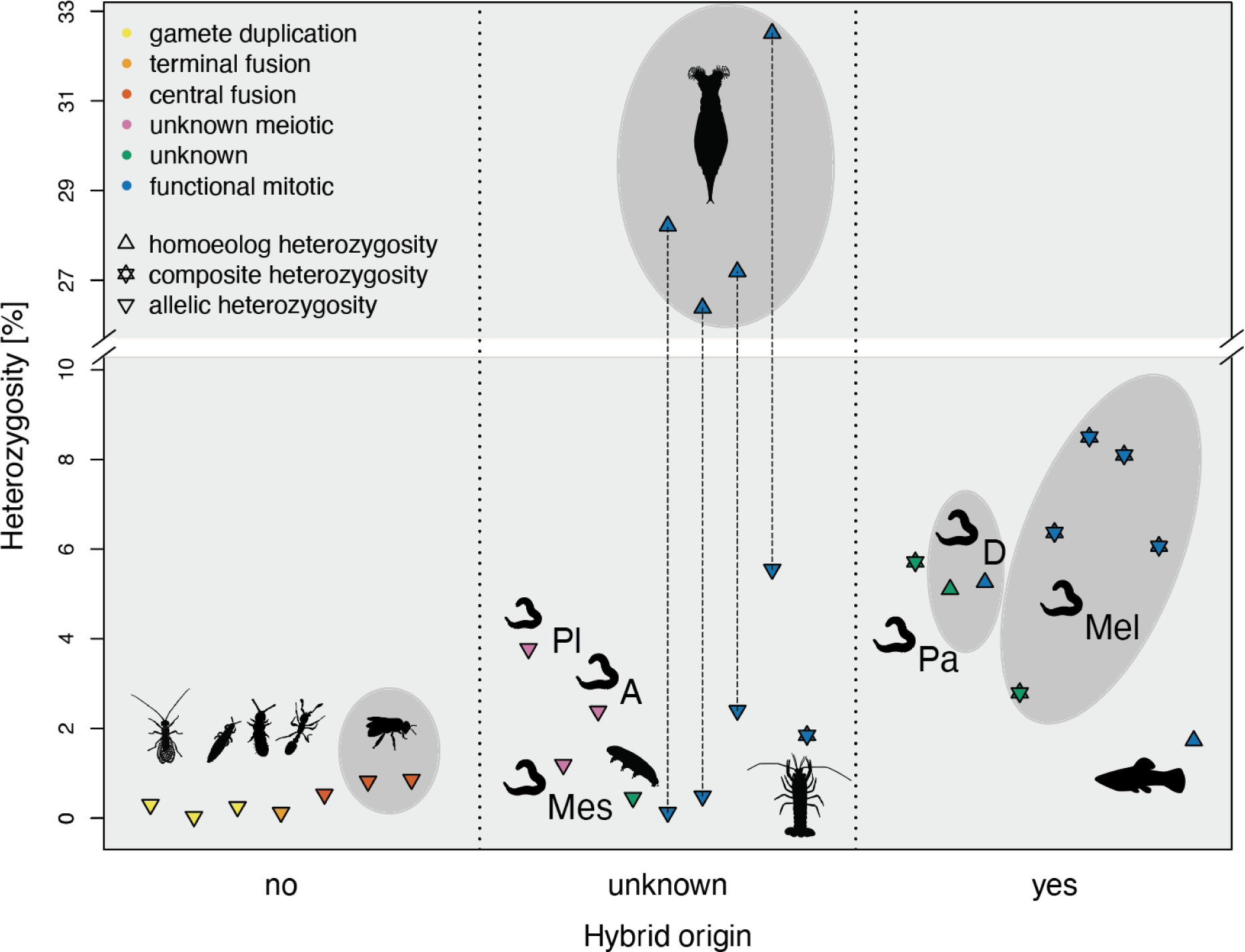
Hybrid origin is the main driver of high heterozygosity in parthenogenetic species. Heterozygosity estimates with respect to hybrid origin (x axis) and cellular mechanism of parthenogenesis (color code). Whenever possible, heterozygosity is decomposed into heterozygosity between alleles of intra-specific origin versus heterozygosity between homoeologs of hybrid origin (shapes). Composite heterozygosity covers the cases where the two cannot be distinguished. Species with a possible shared origin of parthenogenesis are grouped in gray ellipses. Nematode genus abbreviations: Pl: *Plectus*, Mes: *Mesorhabditis*, D: *Diploscapter*, Pa: *Panagrolaimus*, A: *Acrobeloides*, Mel: *Meloidogyne*. The estimates for each species are listed in **Supplementary Table 2**, species names for the silhouettes are indicated in Figure 1.

The heterozygosity levels present at the inception of parthenogenesis should decay over time for most forms of meiotic parthenogenesis [10,59] (see also **Box 2**). In functionally mitotic parthenogens, heterozygosity is expected to increase over time as haplotypes can accumulate mutations independently of each other (generating the so-called ‘Meselson effect’) [4]. However, gene conversion can strongly reduce haplotype divergence and, if high enough, can even result in a net loss of heterozygosity over time, even under functionally mitotic parthenogenesis [4,13]. In spite of the prediction that the cellular mechanism of parthenogenesis should affect heterozygosity, it appears to have no detectable effect on heterozygosity levels once we control for the effect of hybrid origins (**Figure 2**). Indeed, heterozygosity levels in the eleven functionally mitotic parthenogens are high, but all these species are of hybrid origin. Furthermore, nine of the eleven species are polyploid (the diploid species are the amazon molly and the nematode *Diploscapter pachys*). Conversely, all the species with meiotic parthenogenesis are diploid. This is expected given that polyploidy can generate problems during meiosis (reviewed in [41]), but complicates the interpretation of heterozygosity levels among species with different cellular mechanisms of parthenogenesis. Nevertheless, for the species that are not of hybrid origin, it is interesting to note that different forms of meiotic parthenogenesis (including gamete duplication, terminal and central fusion) are associated with similarly low heterozygosity levels. This suggests that although the rate of heterozygosity loss is expected to vary according to mechanisms of parthenogenesis, this variation is only relevant very recently after transitions to parthenogenesis, and no longer affects heterozygosity among established parthenogenetic species. Consistent with this view, the two non-hybrid parthenogens with relatively higher heterozygosity are the youngest ones in our dataset: 40 years for the cape bee [20] and 100 years for the raider ant [19]. Alternatively, variation in heterozygosity caused by different forms of meiotic parthenogenesis may be too small to be detected with our methods.

### Heterozygosity structure in polyploids

In polyploids the estimated genome-wide heterozygosity can be generated by a single genome copy that is highly divergent while others are similar, or by homogeneous divergence across all copies present, or a combination of these. We therefore characterized the most likely origin of heterozygosity for the polyploid species in our dataset. The heterozygosity of two of the five triploid species in our dataset (the crayfish *P. virginalis* and nematode *M. floridensis*) is generated mostly by biallelic loci (**Figure 3, Supplementary Materials S3**). The very low proportion of triallelic loci in these two genomes suggests an AAB structure of the two genomes, where two of the haploid genome copies (A) are nearly identical and the last copy (B) is the carrier of the observed heterozygosity. This AAB model is in agreement with the previous genomic analysis of the parthenogenetic crayfish *P. virginalis* [25]. This previous analysis suggested that *P. virginalis* has a non-hybrid origin, which would imply that the divergence between the A and B haploid genomes corresponds to the heterozygosity present in the sexual ancestor *P. fallax* from which *P. virginalis* derived only 30 years ago. However, our estimation of the divergence of the third genome copy B (1.8%) exceeds by far our estimation of the heterozygosity in *P. fallax* (0.76%; **Supplementary Materials S4**). We therefore suggest a hybrid origin of *P. virginalis*, similar to the well established hybrid origins of the other polyploid asexuals in our dataset. The second triploid species in our dataset with an AAB genome structure is the root knot nematode *M. floridensis*, which was previously mistaken for diploid (**Supplementary Materials S1**). This genome structure distinguishes *M. floridensis* from the other triploid *Meloidogyne* species (whose genomes are comprised of a larger portion with an ABC structure, **Figure 3**), which might suggest that the origin of triploidy in *M. floridensis* is independent of the origin of triploidy in the other *Meloidogyne* species.

**Figure 3:**
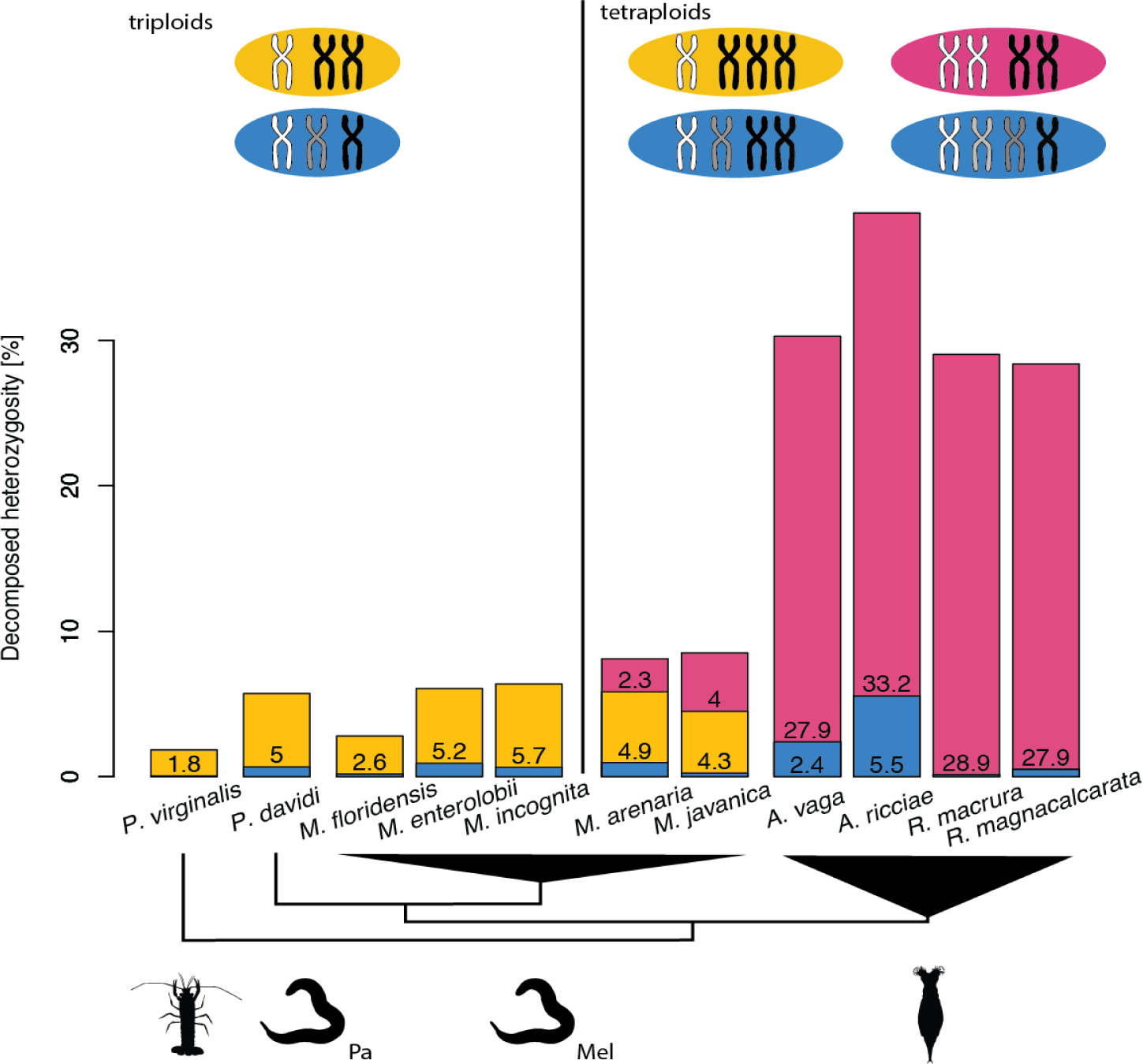
The relative heterozygosity structure in polyploids. Biallelic loci are indicated in yellow or pink: yellow when the alternative allele is carried by a single haplotype (AAB or AAAB), and pink when both alleles are represented twice (AABB). Loci with more than two alleles are indicated in blue.

In the tetraploid species, the biallelic loci can be sorted into those where one genome copy carries the alternative allele (yellow portions in **Figure 3**), and those where two genomic copies carry the alternative allele (pink portions). The genomes of the two tetraploid *Meloidogyne* species contain high proportions of all heterozygosity structures (**Figure 3**) suggesting a complex genomic structure, such as AABC or ABCD. Alternatively, this signal could be also caused by partial aneuploidy, which is common among *Meloidogyne* species [60].

Haplotype divergence can provide information on the origin of parthenogenesis in polyploid species: in asexual polyploids of hybrid origin we expect, and indeed observe, highly heterogeneous divergences among haplotypes, while in those of intra-specific origin we expect homogeneous divergences. Notably, divergence between the haplotypes of the four bdelloid rotifers is highly asymmetric (**Figure 3**). When tetraploidy was first discovered in bdelloids, it was proposed that it stemmed from either a whole genome duplication or a hybridization event in their ancestor [61]. Studies of bdelloid rotifers traditionally refer to the divergent haplotypes as “ohnologs” (e.g., [13,14]), which, following the unified vocabulary of Glover et al [62] would imply that the diverged haplotypes are products of a whole genome duplication. However, the most parsimonious explanation for the highly asymmetric divergence of the different bdelloid haplotypes is a hybrid origin. Referring to the diverged haplotypes as homoeologs therefore appears more appropriate. Our analyses also indicate that the allelic heterozygosity varies extensively among bdelloid rotifer genera. Divergence is very low in *Rotaria* (0.49% in *R. magnacalcarata* and 0.125% *R. macrura*) but relatively high in *Adineta vaga* (2.4%) and in *A. ricciae* (5.5%). There is currently no good explanation for the higher allelic heterozygosity in *A. ricciae* compared to *A. vaga*, but our analyses of the *A. ricciae* sequencing reads suggest that different ploidy levels in the two species could play a role (**Supplementary Materials S2**). Additionally, variable gene conversion and mutation rates could also contribute to the different allelic heterozygosities. In *A. vaga* it has been suggested that gene conversion reduces divergence between homologs in some genome regions [13] but there are currently no direct estimates for gene conversion rates in rotifers. Independently of the mechanisms causing the differences between bdelloids, it is important to note that with such low levels of divergence between homologs, there can be no strong genome-wide ‘Meselson effect’ in bdelloid rotifers (see also [13]). It remains possible that the subset of genomic regions with divergence between homologs in *Adineta* features allele phylogenies as expected under the ‘Meselson effect’. This is the case in the asexual unicellular eukaryote *Trypanosoma brucei gambiense*: some genome regions feature high heterozygosity and allele phylogenies as expected under the ‘Meselson effect’, while others are largely homozygous [63]. Again, it remains unknown why there is such extensive heterogeneity in divergence across the genome in this species. A possible explanation is that the heterozygous genome regions are the consequence of ancient introgression, and that gene conversion rates are low in such regions because of their very high heterozygosity (see **Conclusions**).

### Palindromes and gene conversion

Palindromes are duplicated regions on a single chromosome in reverse orientation. Because of their orientation, palindromes can align and form hairpins, which allows for gene conversion within duplicated regions (**Supplementary Figure 2**). Palindrome-mediated gene conversion was shown to play a major role in limiting the accumulation of deleterious mutations for non-recombining human and chimpanzee Y chromosomes [47,48,64]. Indeed, approximately one third of coding genes on these Y chromosomes occur in palindromes, and the highly concerted evolution of palindromic regions indicates that the rates of gene conversion are at least two orders of magnitude higher in the palindromes than between homologous chromosomes [64]. The reports of palindromes in the genomes of the bdelloid rotifer *A. vaga* [13] and of the springtail *Folsomia candida* [21] led to the hypothesis that palindromes could play a similar role in asexual organisms by reducing deleterious mutation accumulation in the absence of recombination. However, the potential benefit of palindrome-mediated gene conversion depends on the portion of genes in palindromic regions [47]. In addition to identifying palindromes, it is therefore important to also quantify the number of genes affected by palindrome-mediated gene conversion.

We identified 19 palindromes in *A. vaga*, 16 in *F. candida*, and one to four palindromes in eight additional genomes (**Table 1**). Not a single palindrome was detected in the remaining 13 species where the available data allowed us to identify palindromes (See **Methods** and **Supplementary Materials S2** for details). The frequency of palindromes had no phylogenetic signal; for example, although we found 19 palindromes in *A. vaga*, we found no palindromes in the three other bdelloid rotifers (in agreement with [14]). However, it is important to note our analyses assume that there are no systematic scaffolding errors in the published assemblies, meaning that our list of palindromes includes false positives that are generated by mis-assemblies in the published reference genomes.There is also no indication for major rearrangements being present solely in very old asexuals; among the very old asexuals, the non-*A. vaga* rotifers along with the *Diploscapter* nematodes have either zero or only a single palindrome.

**Table 1:**
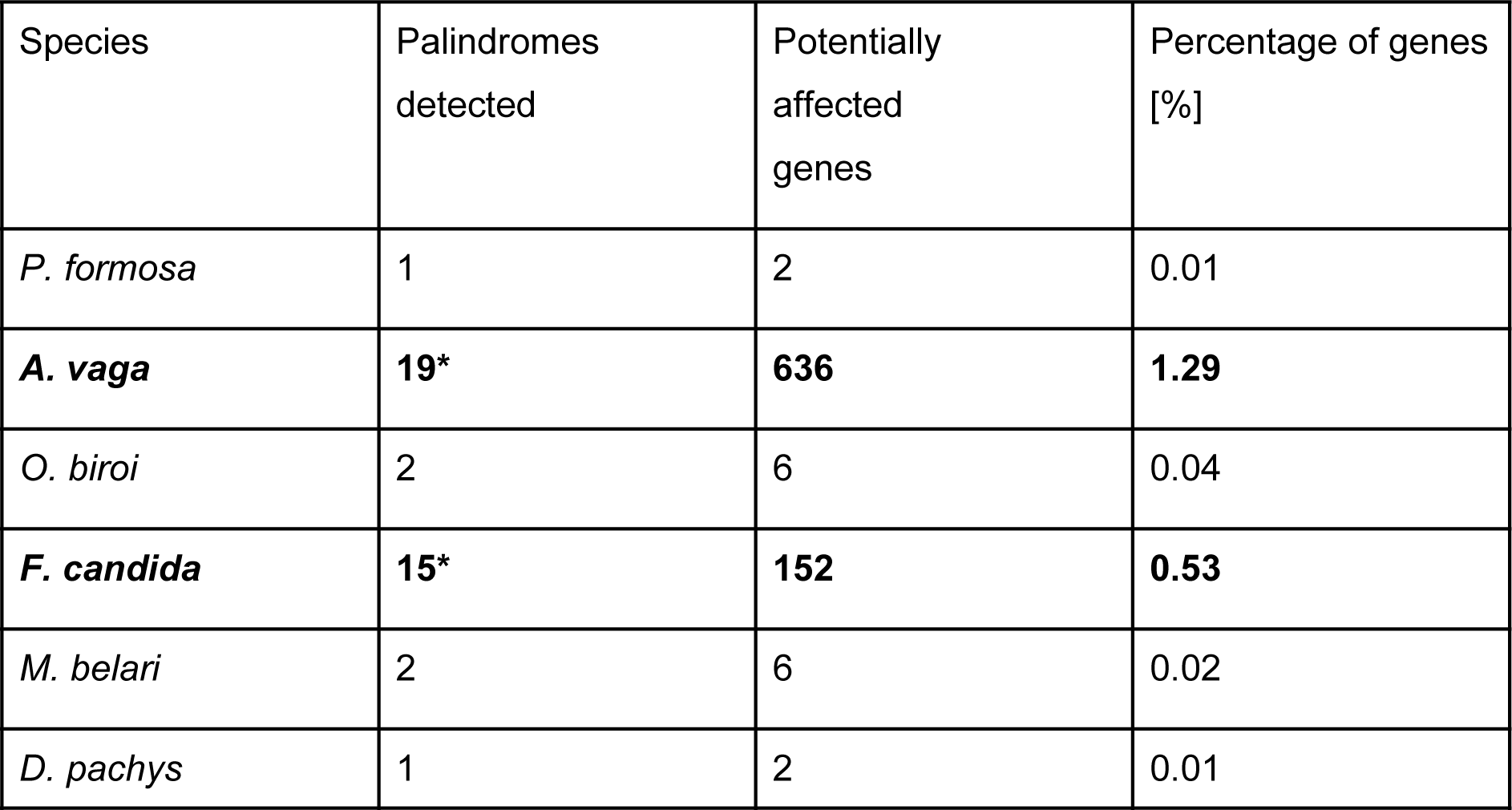
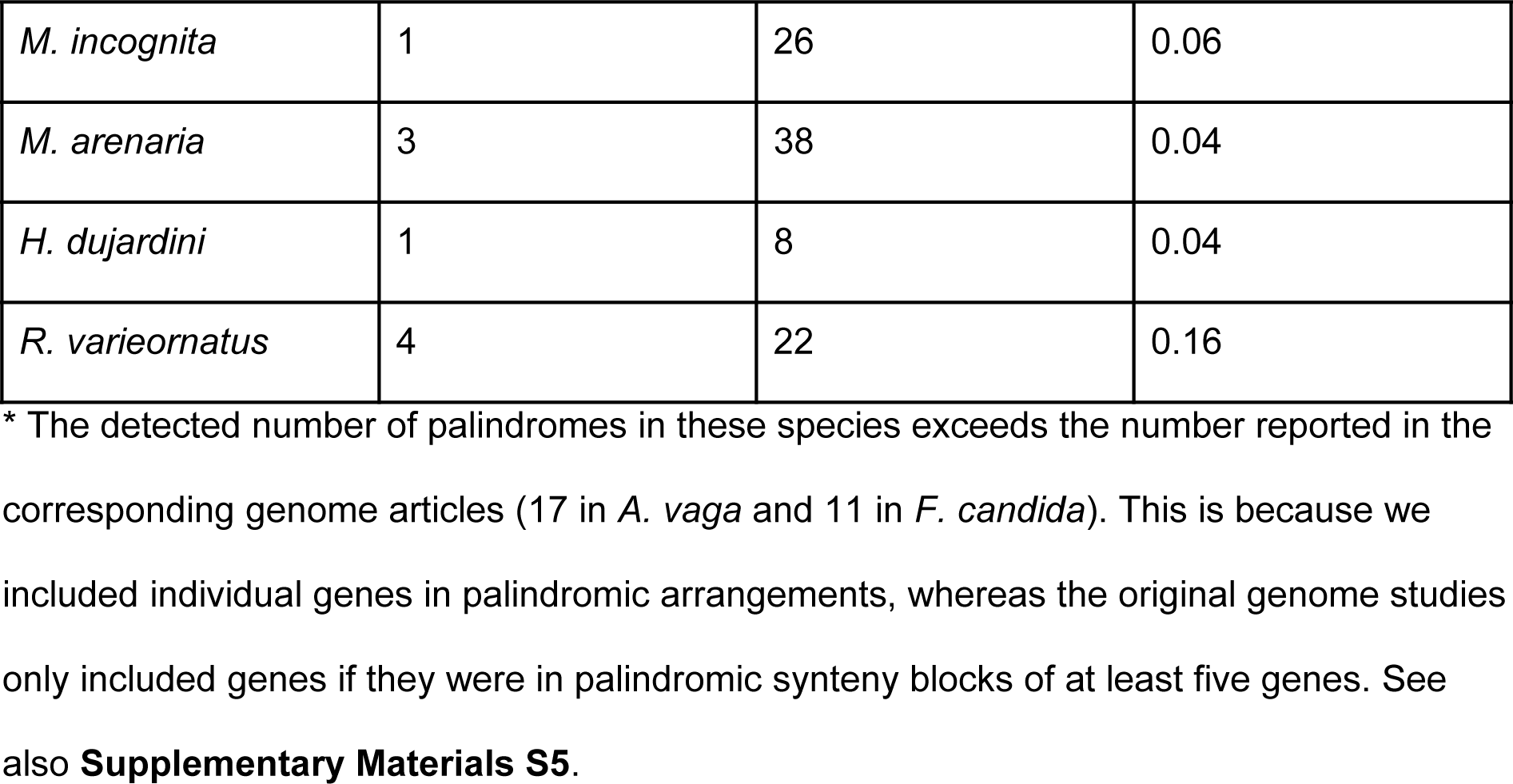
Palindromes in parthenogenetic genomes. Only species with at least one palindrome detected are listed in the table. Rows in bold highlight species with more than 100 genes detected in palindromes.

*Adineta vaga* and *F. candida* are the only two species with more than 100 genes potentially affected by palindrome-mediated gene conversion, but even for these two species, the overall fraction of genes in palindromes is very small (1.23% and 0.53% respectively). The fraction of genes in the other seven species ranges between 0.01% and 0.16%, suggesting that palindromes do not play a major role in the genome evolution of any of the parthenogenetic lineages analyzed. Our findings substantiate the conclusion of a previous study [14] that major genomic rearrangements and the breaking of gene syntenies do not occur at high rates in asexual organisms. They appear to occur at rates similar to those known in recombining genome portions of sexual species [65,66].

Gene conversion can also occur outside of palindromic regions between alleles, for example when double-stranded DNA breaks are repaired using the homologous chromosome as a template [67,68]. This can, in theory, contribute to the loss of heterozygosity under all forms of parthenogenesis, but allelic gene conversion rates have only rarely been studied in parthenogenetic species – or sexual ones for that matter. Allelic gene conversion rates are estimated differently in different studies and are therefore difficult to compare: in the water flea *D. pulex*, they were estimated to amount to approximately 10^−6^ locus^−1^ generation^−1^ [22,23,69], and in the amazon molly *Poecilia formosa* to 10^−8^ [12]. Up to 11% of the genome of the nematode *D. pachys* [27] is suggested to be homozygous as a consequence of gene conversion, and studies have also argued for an important role of gene conversion for genome evolution in root-knot nematodes [32] and rotifers [13,14], although no quantitative estimates are available for these groups.

### Transposable elements

Transposable elements (TEs) are DNA sequences that can autonomously change positions in a genome via various ‘cut-and-paste’ and ‘copy-and-paste’ mechanisms [70,71]. TEs can invade genomes even though they generally provide no adaptive advantage to the individual carrying them [72–74]. To the contrary, new TE insertions in coding or regulatory sequences disrupt gene functions and cause deleterious effects in the host; only very rarely can specific insertions be co-opted to acquire novel, adaptive functions for the host [74]. In sexual organisms, TEs can spread through panmictic populations because of their ability to rapidly colonize new genomes [6,75]. At the same time, sexual reproduction facilitates the purging of deleterious TE insertions, because recombination, segregation, and genetic exchange among individuals improve the efficacy of selection [76,77]. In the absence of sex, TEs could therefore accumulate indefinitely, which led to the prediction that TEs could frequently drive the extinction of parthenogenetic lineages. Only parthenogenetic lineages without active TEs, or with efficient TE suppression mechanisms, would be able to persist over evolutionary timescales [77,78]. Consistent with this view, initial investigations in bdelloid rotifers reported extremely low TE loads [79]. This prompted the authors to suggest that bdelloid rotifers could have been able to persist in the absence of canonical sex for over 40 million years thanks to their largely TE-free genomes.

Our analysis of parthenogenetic animal genomes does not support the view that bdelloid rotifers have unusually low TE contents, even though our methods tend to underestimate TE content in bdelloids relative to other parthenogenetic species (see **Supplementary Methods**). The estimated TE content of bdelloid rotifers (0.7% to 9.1%) is comparable to other parthenogenetic animal taxa (median 6.9%, **Figure 4**), all of which are considerably younger than the bdelloids. Across the 26 genomes, there was large variation in total TE content, overall ranging from 0.7% to 17.9%, with one species, the marbled crayfish, reaching 34.7%. The extreme repeat content in the latter is largely inherited from its sexual ancestor, as the TE loads we estimated for the close sexual relative *P. fallax* are nearly identical (32.5%). Nevertheless, the abundance of TEs in parthenogenetic animal genomes appears to be generally lower than in sexual species, which range typically from 8.5-37.6% (median: 24.3%) [80]. These low abundances could stem from the evolution of reduced TE activity in parthenogenetic species [81,82], and/or appear if successful parthenogenetic taxa generally derive from sexual ancestors with largely inactive TEs and low TE contents. However, whether the apparently lower TE content in the 26 genomes is indeed linked to parthenogenesis remains an open question as TE loads are known to be highly lineage-specific [17,33,83,84].

**Figure 4:**
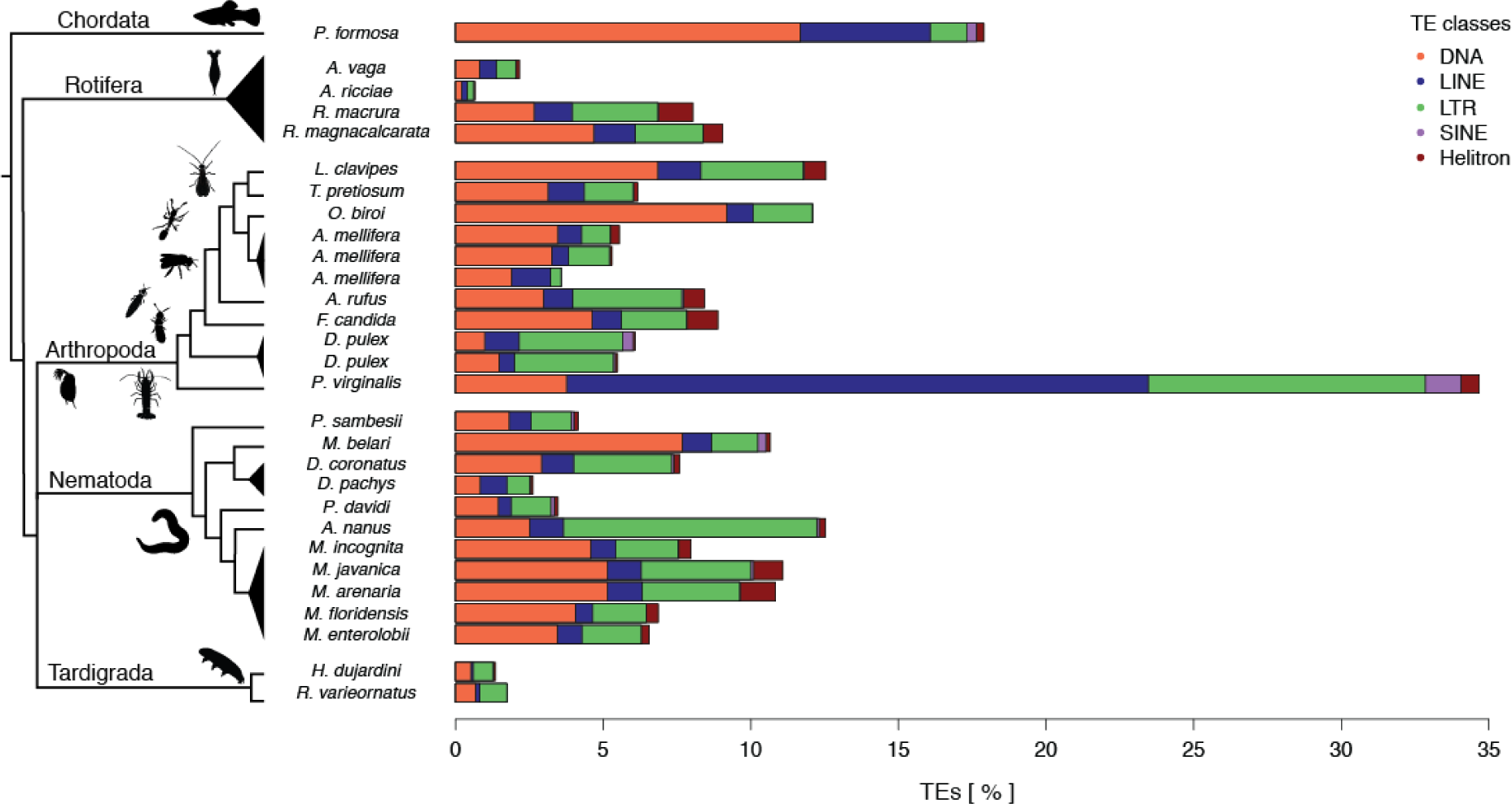
Percentage of nucleotides annotated as transposable elements (TEs) in parthenogenetic genomes. Both the TE load and frequency of TE classes vary substantially between individual parthenogenetic lineages. The TE classes are: class I “cut-and-paste” DNA transposons (DNA), and class II “copy-and-paste” long interspersed nuclear elements or autonomous non-LTR elements (LINEs), short interspersed nuclear elements or non-autonomous non-LTR elements (SINEs), long terminal repeat elements (LTR), and rolling-circle elements (Helitron).

In addition to other lineage-specific characteristics, the cellular mechanisms underlying parthenogenesis could also affect TE loads. For example, many forms of meiotic parthenogenesis can allow for the purging of heterozygous TE-insertions, given the loss of heterozygosity between generations **(Box 2)**. However, in the genomes analyzed here, we did not find any effect of cellular mechanisms on TE loads (**Supplementary Figure 8**), likely because the expected effect of the cellular mechanisms is small relative to lineage-specific mechanisms. Moreover, host TE suppression mechanisms can contribute to the inactivation and subsequent degeneration of TE copies over time, independently of the cellular mechanism of parthenogenesis [74,85]. Similarly, we did not find any difference in TE content according to hybrid *versus* intraspecific origin of asexuals (**Supplementary Figure 8**), even though mismatches between species-specific TEs and silencing machineries can result in increased TE activity in hybrids [7,9,86–88].

### Horizontal gene transfer

Parthenogenetic species could harbour many genes acquired via horizontal gene transfer (HGT) as a consequence of relaxed selection on pairing of homologous chromosomes (see also section **Gene family expansions**). It has also been proposed that HGTs might represent an adaptive benefit which allows for the long-term maintenance of parthenogenesis [89].

We found no evidence for parthenogenesis favoring the retention of HGTs. The majority of the species for which the available data allowed for HGT inferences (16 out of 23) showed a low proportion of candidate HGT genes, with ∼1% candidate HGT (**Supplementary Table 4**). In agreement with previous findings, we identified elevated levels of candidate HGT in the four bdelloid rotifer species *A. ricciae* (10%), *A. vaga* (10.6%), *R. macrura* (8.4%) and *R. magnacalcarata* (7.2%) and the springtail *F. candida* (6.3%). However, we also detected unexpectedly high levels of candidate HGT in the ant *O. biroi* and the wasp *T. pretiosum* (**Supplementary Table 4**). To evaluate a potential link between elevated candidate HGT levels and parthenogenesis in hexapods we additionally quantified candidate HGT in a number of published genomes of sexual species from the same order or superfamily as the parthenogenetic species. This revealed a similarly high proportion of candidate HGT in these sexual relatives, suggesting that whatever the cause of high candidate HGT in these taxa is a general characteristic of hexapods and not linked to the switch to parthenogenetic reproduction. The precise nature of these putative foreign genes in hexapod genomes remains unknown.

### Gene family expansions

Most genome papers scan for expansions of specific gene families. Such expansions are then discussed in the light of the focal species’ biology. The expansion of specific gene families *per se* is thus generally a species-specific trait [90] that is not related to parthenogenesis. To our knowledge, the only example of a gene family expansion that could be directly associated with parthenogenesis is the diversification of the RNA silencing machinery of TEs in bdelloid rotifers [13]. TEs are expected to evolve reduced activity rates in parthenogenetic hosts (see section **Transposable elements**), and an improved RNA silencing machinery could be the mechanism underlying such reduced activity rates.

Functionally mitotic parthenogenesis might facilitate variation in gene copy numbers between homologous chromosomes as a consequence of relaxed constraints on chromosome pairing. Gene family expansions (and contractions) could therefore be more extensive and be retained more frequently in parthenogenetic than sexual species. To test this hypothesis, an overall comparison of gene family expansions in sexual and parthenogenetic sister species is needed (see **Supplementary Materials S5**). Four studies have surveyed gene family expansions in parthenogenetic species as well as in (sometimes distantly related) sexual counterparts, but these studies found no differences between reproductive modes [12,18,21,29]. However, only two of the four studies are based on parthenogens with functionally mitotic parthenogenesis (i.e., where chromosome pairing is not required), and additional studies are therefore needed to address the question of whether parthenogenesis affects gene family expansions.

### Gene loss

Parthenogenetic animals are predicted to lose genes underlying sexual reproduction traits, including male-specific traits and functions (e.g. male-specific organs, spermatogenesis), as well as female traits involved in sexual reproduction (e.g., pheromone production, sperm storage organs) [91]. In the absence of pleiotropic effects, gene loss is expected due to mutation accumulation in the absence of purifying selection maintaining sexual traits, as well as to directional selection to reduce costly sexual traits [92]. Some gene loss consistent with these predictions is documented. For example, the sex determination genes *xol-1* and *tra-2* are missing in the nematode *D. coronatus* [26]. Furthermore, genes believed to be involved in male functions harbour an excess of deleterious mutations in the wasp *Leptopilina clavipes* [16], which could represent the first step towards the loss of these genes. However, a similar excess of deleterious mutations in genes with (presumed) male-specific functions was not detected in the amazon molly *P. formosa* [12].

Parthenogenetic species are further predicted to lose genes specific to some meiotic processes that no longer take the place during egg production [93]. The genes involved in meiosis have been studied in seven parthenogens (the amazon molly, four bdelloid rotifers, and two *Diploscapter* nematodes). There was no apparent loss of meiosis genes in the amazon molly *P. formosa* [12]. The majority of meiosis genes was recovered in bdelloid rotifers [13,14], and furthermore, nearly all of the meiosis genes missing in bdelloid rotifers were also absent in sexual monogonont rotifers, indicating that meiosis gene loss is not associated with the evolution of obligate parthenogenesis [94]. Both *Diploscapter* nematodes also lack certain meiosis genes, but it is unknown whether these genes are also missing in sexual relatives [26,27]. As much as the idea is appealing, there does not seem to be any support for the predicted loss of meiotic genes in functionally mitotic parthenogens. We note that the lack of our understanding of meiosis on the molecular level outside of a few model organisms (particularly yeast and *C. elegans*) makes the interpretation of gene loss (or absence thereof) difficult. This is best illustrated by the fact that losses of presumed “core” meiosis genes have also been reported in different sexual species, where meiosis is clearly fully functional [95].

In summary, some gene loss consistent with the loss of different sexual functions has been reported in several parthenogenetic species. However, there is no striking pattern relative to sexuals, and a clear interpretation of gene loss in parthenogenetic species is problematic because the function of the vast majority of genes is unknown in these non-model organisms.

## Conclusions

We re-analyzed 26 published genomes of parthenogenetic animals to identify genomic features that are characteristic of parthenogenetic animals in general. Many of the original genome studies highlighted one or a few specific features in their focal species, and suggested that they might be linked to parthenogenesis. However, our analyses combined with reviewing published results show that none of these genome features appear to be a general consequence of parthenogenesis, given that none of them was replicated across even a majority of analyzed species.

The variation among genomes of parthenogenetic species is at least in part due to species- or lineage-specific traits. But variation among the features detected in the published single-genome studies is also generated by differences in the methods used. Such differences are often less obvious, yet they can be critical in our assessment of genome diversity among animals. In this work we thus re-analyzed several key genome features with consistent methods. To minimize the potentially confounding effects of differences in assembly quality, we have utilised methods that analyse directly sequencing reads. For example, re-estimating heterozygosity levels directly from reads of each species allowed to show a strong effect of hybrid origin, but not of cellular mechanism of parthenogenesis (**Figure 2**). Another advantage of using the same methods for each species is that it diminishes the “researcher degrees of freedom” [96–98]. For example, the analysis of polyploid genomes requires choosing methods to call heterozygosity and ploidy. By providing a common framework among species, we have shown that homoeolog divergence is very diverse among polyploid asexuals.

We have identified hybrid origin as the major factor affecting heterozygosity levels across all parthenogenetic animal species with available genomic data. This is consistent with the conclusions of two studies that focussed on individual asexual lineages: hybridization between diverse strains explains heterozygosity in *Meloidogyne* root knot nematodes and in *Lineus* ribbon worms [32,99]. This rule applies more generally to all the species analysed with known transitions to parthenogenesis, but it is important to highlight that all the non-hybrid species in our dataset are hexapods. Thus in principle the low heterozygosity could be a hexapod specific pattern, for example due to high mitotic gene conversion rates in hexapods. The taxonomic range of the sequenced species is wide but we are missing several clades rich in parthenogenetic species, such as mites or annelids [100,101]. These clades would be useful foci for future genomic studies of parthenogenetic species.

Independently of the findings of such future studies, our results suggest that heterozygosity loss via meiosis and/or gene conversion plays a significant and highly underappreciated role in the evolution of parthenogenetic species of intraspecific origin and support the theoretical argument that one of the main benefits of sex could be the masking of recessive deleterious mutations (referred to as “complementation”) [102,103]. Conversely, high rates of heterozygosity loss could also allow for the purging of deleterious mutations, as in highly selfing species (e.g. [104,105]). Such purging could help explain why most of the genome scale studies did not find support for the theoretical expectation that parthenogenetic reproduction should result in increased rates of deleterious mutation accumulation (see section **Mutation accumulation and positive selection)**. More generally, given the major differences in genome evolution for parthenogens of intra-specific vs. hybrid origin, our study calls for future theoretical approaches on the maintenance of sex that explicitly consider the loss *vs*. the maintenance of heterozygosity in asexuals.

In our evaluation of the general consequences of parthenogenesis, we were not able to take two key aspects into account: survivorship bias of parthenogenetic lineages, and characteristics of sexual ancestors. How often new parthenogenetic lineages emerge from sexual ancestors is completely unknown, but it has been speculated that in some taxa parthenogenetic lineages might emerge frequently, and then go extinct rapidly because of negative consequences of parthenogenesis. In other words, parthenogens that would exhibit the strongest consequences of parthenogenesis, as predicted by theoretical models, are expected to go extinct the fastest. Such transient parthenogens remain undetected in natural populations, because research focuses on parthenogenetic species or populations, and not on rare parthenogenetic females in sexual populations. Indeed, most of the species included in our study have persisted as parthenogens for hundreds of thousands to millions of years. They might thus be mostly representative of the subset of lineages that suffer weaker consequences of parthenogenesis. Finally, the key constraint for identifying consequences of parthenogenesis is that almost none of the published genome studies of parthenogenetic animals included comparisons to close sexual relatives. This prevents the detection of specific effects of parthenogenesis, controlling for the variation among sexual species – which is extensive for all of the genome features we analyzed and discussed in our study. Overall, despite the importance of recombination rate variation for understanding the evolution of sexual animal genomes (e.g., [106,107]), the genome-wide reduction of recombination does not appear to have the dramatic effects which are expected from classical theoretical models. The reasons for this are probably a combination of lineage-specific patterns, differences according to the origin of parthenogenesis, and survivorship bias of parthenogenetic lineages.

## Data Availability

The code of the pipeline for gathering data, calculating and plotting results is available at https://github.com/KamilSJaron/genomic-features-of-parthenogenetic-animals and it’ s archived in Zenodo (DOI: <u>10.5281/zenodo.3897309</u>); the majority of sequencing reads are available in public databases under the accessions listed in **Supplementary Table 1** The data without publicly available sequencing reads were obtained via personal communication with the corresponding authors.

## Abbreviations

TE: transposable element;
HGT: horizontal gene transfer

## Acknowledgments

We would like to thank Deborah Charlesworth, Marie Delattre, Daniel Wegmann, Christelle Fraisse, Ken Kraaijeveld, Philipp H. Schiffer, Jean-Francois Flot, Dave Lunt, Yuko Ulrich, Sean McKenzie, Julian Gutekunst, Mark Blaxter, Andrés G. de la Filia and anonymous reviewers for comments and discussions. We would also like to thank all the authors who deposited their data in public databases with correct metadata and all the authors who were helpful when we asked for data or clarifications.

## Author contributions

K.S.J. and T.S. conceived the study, K.S.J. and J.B. collected the data; K.S.J., R.W.N. and T.R.R.B. performed the analyses; K.S.J., J.B. and T.S. reviewed the literature; K.S.J., T.S., J.B., M.R.R. and R.W.N. wrote the manuscript; All authors were involved in discussions about results and interpretations.

