## Supplementary table 1 for "Genomic features of parthenogenetic animals"

**Supplementary table 1: Overview of analysed species.** This information was collected directly from the cited literature. References include information regarding cellular mode of reproduction, origin of asexuality and/or the age of asexuality.

| species | common name | NCBI accesion | cellular mechanism of parthenogenesis | evidence | hybrid origin | age of asexuality [y] | references |
|---|---|---|---|---|---|---|---|
| *Poecilia formosa* | amazon molly | SAMN01797685 | sperm-dependent functional mitotic | cytology, genetics | yes | 100 k | [1] |
| *Adineta vaga* | bdelloid rotifer | SAMEA2043852 | functional mitotic | stained karyotypes have no apparent chromosome pairs | | 46 M | [2, 3] |
| *Adineta ricciae* | bdelloid rotifer | SAMEA104393659 | functional mitotic | | | 46 M | [3, 4] |
| *Rotaria macrura* | bdelloid rotifer | SAMEA104393678 | functional mitotic | no direct evidence[1] | | 46 M | [5, 6] |
| *Rotaria magnacalcarata* | bdelloid rotifer | SAMEA104393684 | functional mitotic | no direct evidence[1] | | 46 M | [5, 6] |
| *Leptopilina clavipes* | parasitoid wasp | SAMN02047179 | gamete duplication[2] | cytology | no | 6-43 k | [7] |
| *Trichogramma pretiosum* | trichogramma wasp | SAMN02439301 | gamete duplication[2] | cytology, genetic markers | no | "few" | [8, 9] |
| *Ooceraea biroi*[3] | raider ant | SAMN02428046 | central fusion | cytology, RAD seq | no | | [10] |
| *Apis mellifera capensis* | cape honey bee | SAMN10245904 SAMN10245906 | central fusion | cytology | no | 20 | [11] |
| *Aptinothrips rufus* | thrip | | gamete duplication[4] | | no | 150-200 k | [12] |
| *Folsomia candida* | springtail | SAMN04196550 | terminal fusion[4] | cytology | no | | [13] |
| *Daphnia pulex* | water flea | SAMN03964753 SAMN03964750 | central fusion equivalent | No separation at meiosis I; abortive meiosis | yes | 1-170 k | [14] |
| *Procambarus virginalis* | marbled crayfish | SAMN07142640 | functional mitotic | microsat study, histological evidence | | less than 30 | [15–17] |
| *Plectus sambesii* | nemotode | SAMN07227113 | unknown meiotic | 2 meiotic divisions, 2 polar bodies, but no fusions were observed, putative endoduplication | | | [18] |
| *Mesorhabditis belari* | nemotode | SAMEA5150020 | unknown meiotic | cytology; 2 meiotic divisions observed | | | [19] |
| *Diploscapter coronatus* | nemotode | SAMD00025087 | unknown | | yes | | [18, 20] |

[1] Meiosis was not observed in two bdelloid rotifers *Habrotrocha tridens* and *Philodina roseola*, both members of *Philodinidae*, the same family as *Rotaria*.
[2] *Wolbachia* induced
[3] formerly *Cerapachys biroi*
[4] suggested that it is endosymbiont induced

| | | | | | | | |
|---|---|---|---|---|---|---|---|
| *Diploscapter pachys* | nematode | SAMN03456257 | functional mitotic | formally central fusion (meiosis I skipped), no recombination, only sister chromatid separation | yes | 18 M[5] | [21] |
| *Panagrolaimus davidi* | nemotode | SAMN02741088 | unknown | polar body produced | yes | 1.3-8.5 M | [22] |
| *Acrobeloides nanus* | nemotode | SAMN06041019 | unknown meiotic | | | | [18] |
| *Meloidogyne incognita* | root-knot nematode | SAMEA104032784 SAMN05712521 | functional mitotic | | yes | ”recently” | [23–25] |
| *Meloidogyne javanica* | root-knot nematode | SAMEA3298191 SAMN05712519 | functional mitotic | | yes | ”recently” | [25] |
| *Meloidogyne arenaria* | root-knot nematode | SAMEA3298190 SAMN05712513 SAMN08721831 | functional mitotic | | yes | ”recently” | [25] |
| *Meloidogyne floridensis* | peach root-knot nematode | SAMN05712529 | unknown | meiotic mechanism suggested by cytology; however the study conflicts in ploidy with the data in this study (see S1) | yes | ”recently” | [25, 26] |
| *Meloidogyne enterolobii* | root-knot nematode | SAMN05712528 | functional mitotic | | yes | ”recently” | [25] |
| *Hypsibius dujardini* | tardigrade; water bear | SAMEA3679301 | terminal fusion equivalent | meiosis II suppressed | | | [27] |
| *Ramazzottius varieornatus* | tardigrade; water bear; Kumamushi | SAMD00054187 | | no males have been found | | | [6] |

[5] Assuming non-hybrid origin suggested by Hiraki at al. 2017.
[6] personal communication with Mark Blaxter
