## Supplementary texts and figures for "Genomic features of parthenogenetic animals"

### Supplementary materials

#### Supplementary Methods

Core genome features (ploidy, haploid genome size, heterozygosity, repeat content, and characterisation of TE content) were estimated directly from sequencing reads to avoid potential assembly biases in reference genome-based approaches. The raw reads were downloaded from SRA using sample accessions specified in **Supplementary Table 1.** The pipeline consisted of several analyses charted on the scheme in **Supplementary Figure 1** and detailed in sections below.

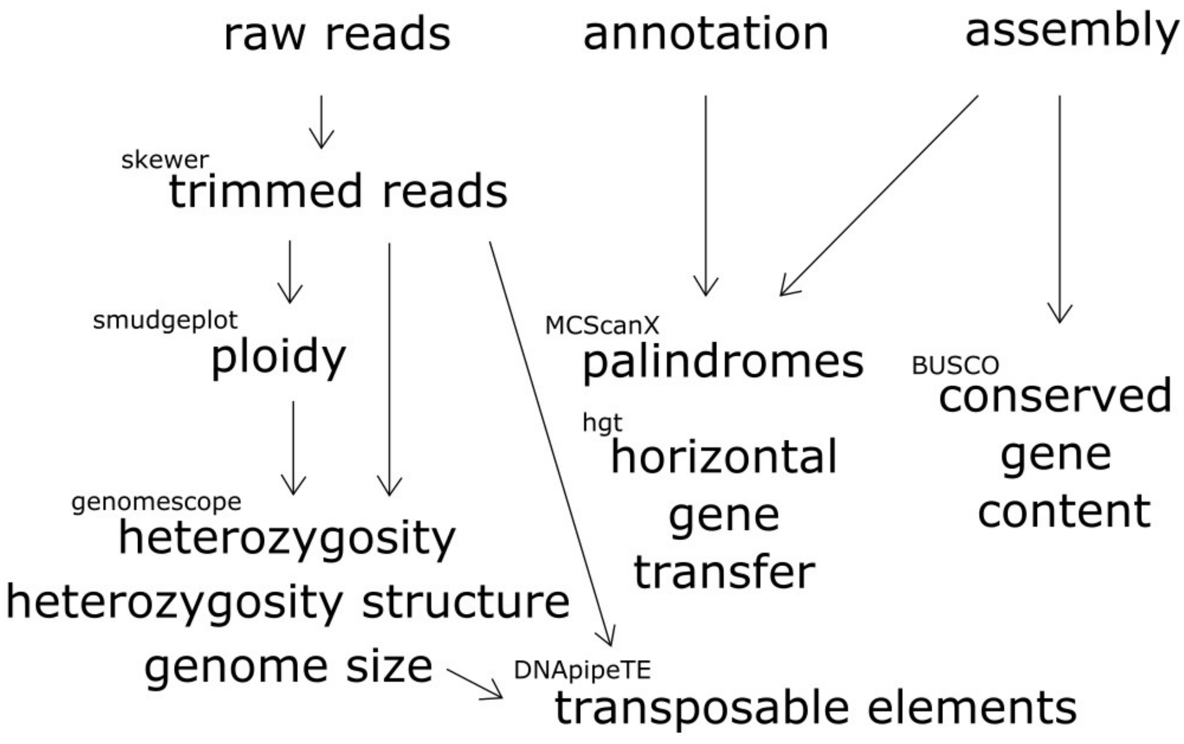


**Supplementary Figure 1:** Scheme of the pipeline**.** Software names used for estimation of individual genomic features are mentioned in a smaller font.

**Raw reads preprocessing**

We cleaned the raw reads by removing standard illumina adaptors (AGATCGGAAGAGCACACGTCTGAACTCCAGTCAC, AGATCGGAAGAGCGTCGTGTAGGGAAAGAGTGTA) and low quality bases (PHRED score < 26) from the 3' end. Only reads longer than 21 bases were retained. The read preprocessing was performed using Skewer (parameters "-z -m pe -n -q 26 -l 21"). [108].

**Genome profiling: ploidy, genome size and heterozygosity**

We used smudgeplot v0.1.3 to estimate ploidy levels [11]. This method extracts from the read set unique kmer pairs that differ by one SNP from each other. These kmer pairs are inferred to derive from heterozygous genome regions. The sum of coverages of the kmer pairs is then compared against their coverage ratio. This comparison separates different haplotype structures (**Supplementary Figure 4b**). The most prevalent structure is then indicative of the overall ploidy of the genome. We used this ploidy estimate in all species, except *A. vaga*. The most prevalent structure suggested that this species is diploid. *A. vaga* is well- characterized as tetraploid [61], but we were unable to detect tetraploidy because homoeologs are too diverged to be identified as such by the kmer-based smudgeplot method.

The definition of heterozygosity for polyploids is not well established, but GenomeScope 2.0 estimates heterozygosity as the proportion of sites that differ in at least one of the homologous regions. This means that in polyploids the estimated genome-wide heterozygosity could be generated by a single haplotype that is highly divergent while others are similar, or by homogeneous divergence across all copies present, or a combination of these.

Using the inferred ploidy levels, we then estimated genome size and heterozygosity using GenomeScope 2.0 [11]. GenomeScope estimates genome wide heterozygosity via kmer spectra analysis, by directly analyzing kmers within the raw sequencing reads. A mixture model of evenly spaced negative binomial distributions is fit to the kmer spectrum, where the number of fitted distributions is determined by the input ploidy. Each distribution corresponds to kmers that occur a given time (e.g. once, twice, etc.) in the genome. Fits are then used to estimate heterozygosity, the fraction of repeats in the genome, as well as the 1n sequencing coverage. The latter is subsequently used for estimation of genome size. The definition of heterozygosity for polyploids is not well established, but GenomeScope 2.0 distinguishes different types of heterozygous loci in polyploids (as shown in **Figure 3**). Specifically, GenomeScope 2.0 utilizes a combinatorial mathematical model to account for how particular nucleotide haplotype structures are related to kmer haplotype structures. Assuming that mutations are randomly distributed across the genome, three equidistant haplotypes will generate the highest fraction of triallelic loci. Conversely, if the divergence is carried by the divergence of a single haplotype, very few or no triallelic loci will be detected.

Kmer spectra analysis is affected by the choice of kmer length. Longer kmers require higher sequencing coverage, but lead to more informative kmer spectra. We have chosen the default kmer size 21 nt for all species except the marbled crayfish, where we chose kmer length 17 nt due to low sequencing coverage.

We were unable to generate heterozygosity estimates for two of the 26 parthenogenetic species for different reasons: in the tardigrade *H. dujardini* because of extensive contamination in the sequencing reads, and in the water flea *Daphnia pulex* samples because of too low coverage.

**Transposable elements**

We quantified transposable elements using DnaPipeTE v1.2 [46]. The method uses the haploid genome size (parameter -genome_size) to subsample sequencing reads to a low coverage of 0.5x (parameter -genome_coverage). We merged all trimmed sequencing reads available for each species for the input of this analysis. These subsampled reads are then assembled using an assembler (Trinity) that can deal with uneven coverages and is able to split assembled regions with few differences (including different TE families). The assembled sequences largely correspond to repetitions as non-repetitive genome regions present in the subsampled reads drop out at this stage, because the coverage of such regions is too low for assembling. The assembled sequences are annotated by homology using a database of known TEs (repbase). This subsampling process is repeated three times (parameter -sample_number), and the union of results represents the repeat library. The third sampling round is used to map overrepresented reads back to the identified TE library to calculate the overall TE abundance based on the fraction of reads mapping to TEs (for details see [46]). The output of the method is the number of sampled nucleotides assembled and annotated as different types of repeats and fractions are calculated as the numbers divided by the total number of sampled nucleotides. Our reported values of TE loads include only repeats that were annotated as TEs, i.e., we did not include 'unknown' repeats which consist of tandem repeats (satellite repeats), duplications or very divergent/ unknown TEs.

We estimated the TE loads in each genome via assembly of overrepresented (i.e., repetitive) genomic sequences and subsequent annotation via homology searches in general databases (see methods). This can result in an underestimation of TE loads in species with many TE types with very low copy numbers (as is the case in *A. ricciae* and

*A. vaga* [13–15]) and in phylogenetically isolated lineages such as rotifers and tardigrades which may comprise divergent TEs not represented in general databases. However, this is unlikely to be the sole reason behind the low TE content of parthenogens reported in our study since the methods we used allowed us to identify similar or higher TE loads in all species without manually curated TE libraries (**Figures 1 and 4**). Manual curation (available for *D. pulex* and *A. vaga* among the species in **Figure 1**) generally increases TE load estimations. For example, manual curation of TEs in the *Daphnia* genome increased estimated TE loads from approximately 9.6% to 17-27% [109]. In *A. vaga*, the recent discovery of a new TE family during manual curation increased the latest TE load estimate to nearly 4% [15].

**Palindromes**

The palindrome analysis was based on genome assemblies and their published annotations. Note that this analysis could not be done for *D. pulex, A. mellifera capensis,* and *A. rufus* because annotations were not available. We performed collinearity analysis using MCScanX (untagged version released 28.3.2013) [49], allowing even a single gene to form a “collinear bloc” (parameter -s) if there were fewer than 100 genes in between (parameter -m). The output was then filtered to contain only blocs on the same scaffold in reverse order. Furthermore we filtered all homologous gene pairs that have appeared on the same strand. All the remaining blocks are palindromes, blocks built of reverse complementary genes on the same scaffold.

We note that it is important to check consistency between the biological interpretation of results, and the methods used to infer them. The default parameters of the software define a collinear block as a sequence of at least 5 genes that are no more than 25 genes apart from each other and then search for such blocks with palindromic arrangement. These default parameters were used in the genome studies of parthenogenetic species (personal communication of the authors of [13,21]). These parameters are geared towards detecting large repeated blocks with large gaps. We argue that small blocks (as small as one gene), but with no gaps within the inverted repeat may also generate gene conversion. Thus, we have reanalysed the genomes allowing for short palindromes of a single gene, because a palindrome could carry fewer than five genes and still be biologically relevant. Re-screening the published genomes for palindromes allowed us to provide a more robust and unbiased view of the importance of palindromes for the evolution of parthenogenetic species.

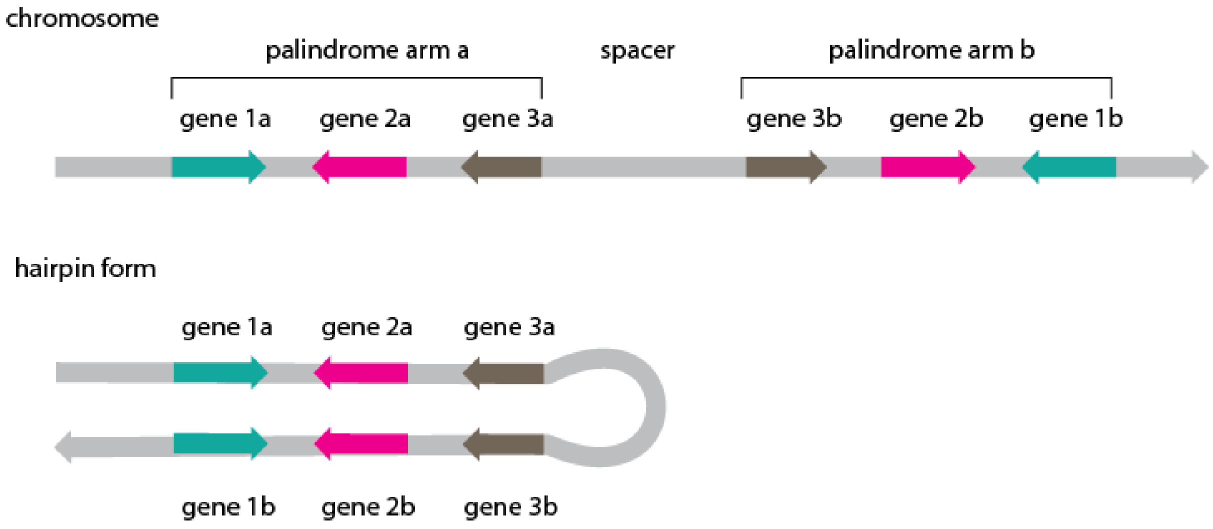


**Supplementary Figure 2: Palindrome structure.** The two homologous reverse complementary regions (arms) of a palindrome are located on the same chromosome. This organisation allows for the formation of a hairpin and can facilitate gene conversion between the palindrome arms.

##### Horizontal Gene Transfer

We assessed the impact of HGT on each parthenogenetic genome using a sequence comparison based approach, following [14]. For each species, the published set of predicted proteins were aligned to the UniRef90 (analysis presented in the main text) and UniProtKB/Swiss-Prot protein databases downloaded on 04/07/2019 and 14/08/2019, respectively [50,110]. The alignment was performed using DIAMOND “blastp” v0.9.21 [111] (“--sensitive -k 500 -e 1e-10”). For each protein, the HGT Index ($h_U$) was then calculated as $h_U = B_{OUT} - B_{IN}$, where $B_{OUT}$ is the bitscore of the best hit to a protein of non-metazoan origin within UniRef90 and $B_{IN}$ is the bitscore of the best hit to a metazoan protein [112]. The proportion of secondary hits that agreed with the designation (metazoan vs non-metazoan) was also recorded as the “consensus hit support” (CHS) [14,34]. To account for the confounding effects of database entries from closely related species contributing to $h_U$, hits from taxa within the same phylum as the focal taxon were excluded from further analysis. Analyses were also run excluding such hits at the class, order and family level for each species, to test the robustness of the results given this partitioning of the target databases (**Supplementary Materials S6**). In each case, a focal protein was designated as a candidate HGT if $h_U > 30$, $CHS_{OUT} > 90\%$, and the protein was found on a scaffold that also encoded at least one gene of unambiguous metazoan origin (i.e., $h_U < 30$ and $CHS_{IN} > 90\%$).

We also estimated HGT candidates in five sexual hexapod species, using the following assemblies: wasps, *Nasonia vitripennis* (GCF_000002325.3) and *Copidosoma floridanum* (GCF_000648655.2); ants, *Camponotus floridanus* (GCF_003227725.1), *Harpagoxenus saltator* (GCF_003227715.1) and *Entomobryomorpha* springtails: *Orchesella cincta* (GCA_001718145.1).

To ascertain the robustness of the detected high levels of $HGT_C$ in the seven species presented in the main text, we also computed $h_U$ based on alignments to the manually curated UniProtKB/Swiss-Prot (i.e., highly conservative) database. The proportion of $HGT_C$

based on comparisons to UniProtKB/Swiss-Prot was substantially lower in *F. candida* (3.13%), *O. biroi* (0.72%) and *T. pretiosum* (2.31%) than that for UniRef90 (but higher for some other taxa, see **Supplementary Table S4**), highlighting the sensitivity of such analyses on the sequence databases used. The proportion of $HGT_C$ observed in bdelloid rotifers also dropped, relative to comparisons against UniRef (2.51 - 3.86%), but remains high relative to other taxa. The impact of the specific database used for HGT detection may be driven by potential taxonomic misclassifications and/or contamination in one or both public databases, or biases in the representative taxa contained within particular databases (e.g., UniProtKB/Swiss-Prot is biased towards a relatively small number of model species). A full investigation of the true composition of public sequence repositories is a major piece of work that is beyond the scope of the current work.

Overall our analysis showed that high levels of HGT are not a general feature linked to parthenogenesis but a clade specific trait of bdelloid rotifers and perhaps of hexapods. However, it is important to note that our analysis does not quantify absolute levels of HGT in any of the taxa; instead, we show the dependency of such estimates on the chosen methodology.

| | mutation accumulation | adaptive evolution | heterozygosity [%] | palindromes [#] | gene conversion [e/(g*nt)] | transposable elements [%] | horizontal gene transfer [%] | gene family expansions | gene loss [l/s] |
|---|---|---|---|---|---|---|---|---|---|
| *P. formosa* | inconclusive [16] | no [16] | 1.43 [16] | | 3e−08 [16] | 16.4 [16] | | no [16] | 0/112 [16] |
| *A. vaga* | | | 1.42 [17, 18] | 17 [17, 18] | [17, 18] | 4 [19] | 7.6 [17, 18] | yes [17] | 4/48 [17, 18] |
| *A. ricciae* | | | 4.55 [18] | 0 [18] | [18] | 2 [18] | 9.1 [18] | | 3/42 [18] |
| *R. macrura* | | | 0.033 [18] | | [18] | 2.2 [18] | 6.2 [18] | | 2/42 [18] |
| *R. magnacalcarata* | | | 0.075 [18] | | [18] | 3 [18] | 6.5 [18] | | 1/42 [18] |
| *L. clavipes* | no [20] | | | | | [21] | | | 16/? [20] |
| *T. pretiosum* | no [22] | | | | | 8.5 [22] | | no [22] | [22] |
| *O. biroi* | | | [23] | | | 4.9 [23] | | | |
| *A. mellifera capensis* | | | [24] | | | | | | |
| *A. rufus* | | | | | | | | | |
| *F. candida* | | | | 11 [25] | | 6.3 [25] | 2.8 [25] | no [25] | |
| *D. pulex* | inconclusive [26] | | 1.62 [26] | | 3.3e−05 [27, 26] | 22.6, 22.0 [21, 28] | | | |
| *P. virginalis* | | | 0.53 [29] | | | 4 [29] | | | |
| *P. sambesii* | | | | | | | | | |
| *M. belari* | | | | | | | | | |
| *D. coronatus* | | no [30] | 5.7 [30] | | | 0.05 [30] | | | 28/96 [30] |
| *D. pachys* | | | 4 [31] | | [31] | | | | 25/91 [31] |
| *P. davidi* | | yes [32] | | | | | 0.63, 0.66 [32] | | |
| *A. nanus* | | | | | | 7 [33] | | no [33] | |
| *M. incognita* | | | 8.4, 3.0 [34, 35, 36] | 1 [35] | [36] | 50, 12 [34, 37, 35] | [34] | | |
| *M. javanica* | | | 7.5, 3.0 [35, 36] | 3 [35] | [36] | 50.8 [35] | | | |
| *M. arenaria* | | | 8.2, 3.0 [35, 36] | 0 [35] | [36] | 50.8 [35] | | | |
| *M. floridensis* | | | 3 [36] | | [36] | 5 [37] | | | |
| *M. enterolobii* | | | [36] | | [36] | | | | |
| *H. dujardini* | | | | | | 15 [37] | 0.8 [38] | | |
| *R. varieornatus* | | | | | | 0.6 [18] | 0.97 [39, 18] | | |

**Supplementary Figure 3: Genomic features studied in parthenogenetic genomes.** The figure mirrors the data from Figure 1, but with references to specific studies added (numbers correspond to the references cited in the main text).

#### S1 Ploidy and reproductive mode of *M. floridensis*

The nematode *M. floridensis* was reported as a diploid species with a mechanism of parthenogenesis functionally equivalent to terminal fusion (absence of the 2nd meiotic division), based on cytological analyses by Handoo et al. [113]. Our analyses indicate that *M. floridensis* is triploid rather than diploid (**Supplementary Figure 4**). Furthermore the heterozygosity detected in this and previous studies [113] is inconsistent with classical terminal fusion (which should result in largely homozygous genomes, see Box 2 and **Figure 2**). Terminal fusion can be associated with high heterozygosity under inverted meiosis (which has been suggested for nematodes of the genus *Acrobeloides* [114]). However, inverted meiosis in *M. floridensis* is rather unlikely given that all other meiotic species in the genus have regular meiosis. We therefore believe that the study of Handoo et al. [113] is either based on an unusual *M. floridensis* strain that has not been used in any genome study thus far or that the cytology inferred by Handoo et al is not correct. These interpretations are further supported by the fact that Handoo et al report on analyses of large numbers of males of *M. floridensis*, while males are unknown/unusual for the strains used in the genome studies. Unfortunately, it is impossible to evaluate the evidence that supported diploidy and terminal fusion in *M. floridensis* as the study by Handoo et al does not include images of chromosomes and egg cells at the basis of their conclusions. Given the genomic evidence is very clear, we consider *M. floridensis* to be triploid for all our analyses and the cellular mechanism of parthenogenesis as “unknown”.

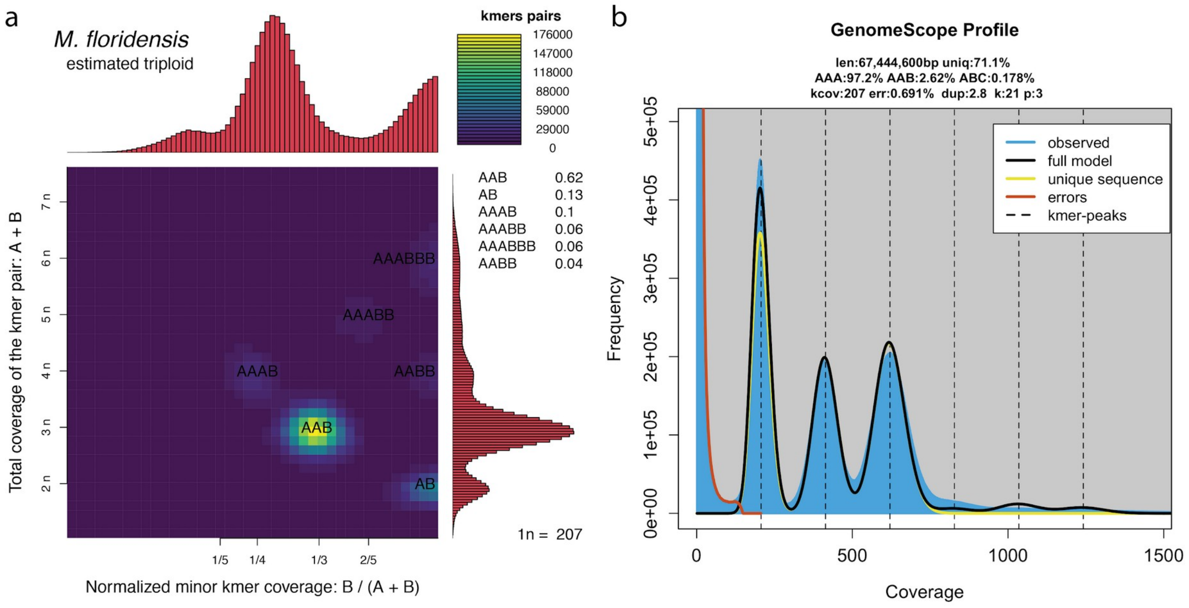


**Supplementary Figure 4: Genomic evidence of triploidy in *M. floridensis*. a |** the smudgeplot shows dominance of a triploid (AAB) genome structure. The smudges corresponding to higher ploïdies are likely originating from paralogs. The diploid kmer pairs (AB) represent situations where the third allele is diverged from the two more than one nucleotide. **b |** kmer spectra analysis of *M. floridensis* shows a typical triploid genome structure with haploid, diploid and triploid peaks and expected distances from each other.

#### S2 Haplotype structure of *Adineta ricciae*

The diploid genome assembly (i.e., where haplotypes are largely separated and present in the assembly) of *A. ricciae* spans 201 Mbp and carries approximately 63,000 genes [14]. Analyses of assembled genome sections suggested a tetraploid genome structure AABB similar to other bdelloid rotifers [14,61]. However, our reanalyses of the *A. ricciae* sequencing reads with kmer spectra appear to be better consistent with an octoploid structure, rather than tetraploid. The interpretation of kmer spectra in rotifers is complicated by the fact that the A and B genomes are very highly diverged (in *A. ricciae*, the average divergence between A and B is 33.21%, while the divergence within AA and BB is 5.55%; **Figure 2**). Given this extreme divergence, A and B share practically no kmers and therefore the kmer spectrum is expected to miss the homozygous peak at 4n and therefore resemble the spectrum of a diploid species (with high heterozygosity). However, if *A. ricciae* was indeed characterized by a tetraploid AABB genome structure, the kmer spectrum should show a big haploid (n) peak at 124x coverage (as calculated from the genome size and total sequencing reads), generated from variation within A and B haplotypes. A second (2n) peak at ~248x would then be generated by homozygous regions within A and B. However, the kmer spectrum of *A. ricciae* does not follow these predictions (**Supplementary Figure 5a**). Instead, we observe three peaks: a haploid peak at 71x coverage (partially overlapping with variation generated by sequencing error), a diploid peak at 142x and a third, tetraploid peak at ~284x (**Supplementary Figure 5b**). We verified that the 1n peak indeed contains complementary kmers and therefore is not generated by sequencing errors or contamination (**Supplementary Figure 5c**). As a second line of evidence, we also examined if the kmers from the 1n peak were present in the decontaminated assembly of *A. ricciae* using KAT [115]. Indeed, the decontaminated assembly consisted of many of the kmers from the 1n peak (**Supplementary Figure 5d**).

If it is the case that homoeologous kmers are entirely excluded from the kmer analyses, the resultant spectrum with three peaks (n=71x, 2n~142x and 4n~284x) is incompatible with the proposed degenerate tetraploid structure based on the current interpretation of the genome assembly [19]. Rather, these patterns are better explained by a 2x higher ploidy level, i.e., to octoploidy. A simple doubling in chromosome number in *A. ricciae* is not supported by the observation of 12 chromosomes in this species, the same as for *A. vaga* [116], and thus any increase in ploidy level in *A. ricciae* would seem to be either complex or very recent. Alternatively, the kmer peak at ~284x might be caused by collapse across all four subgenomes, creating regions of the genome with 0% divergence across both homologous and homoeologous copies. However, this hypothesis is at odds with observations of read- and SNP-coverage from [19], that indicate the majority of the diploid assembly (~81%) is twofold covered, consistent with the idea of large-scale increases in ploidy level. Finally, it is not entirely precluded that the raw data itself, or the methods used to generate it, contains some stratification that might produce a false signal of increased ploidy. Further investigation of these alternative hypotheses will likely require additional data, such as long-read sequencing technology, which is beyond the scope of the present study. However, the likely differences in ploidy levels between *A. vaga* and *A. ricciae* preclude a clear interpretation of the strikingly different allele divergences between these two species.

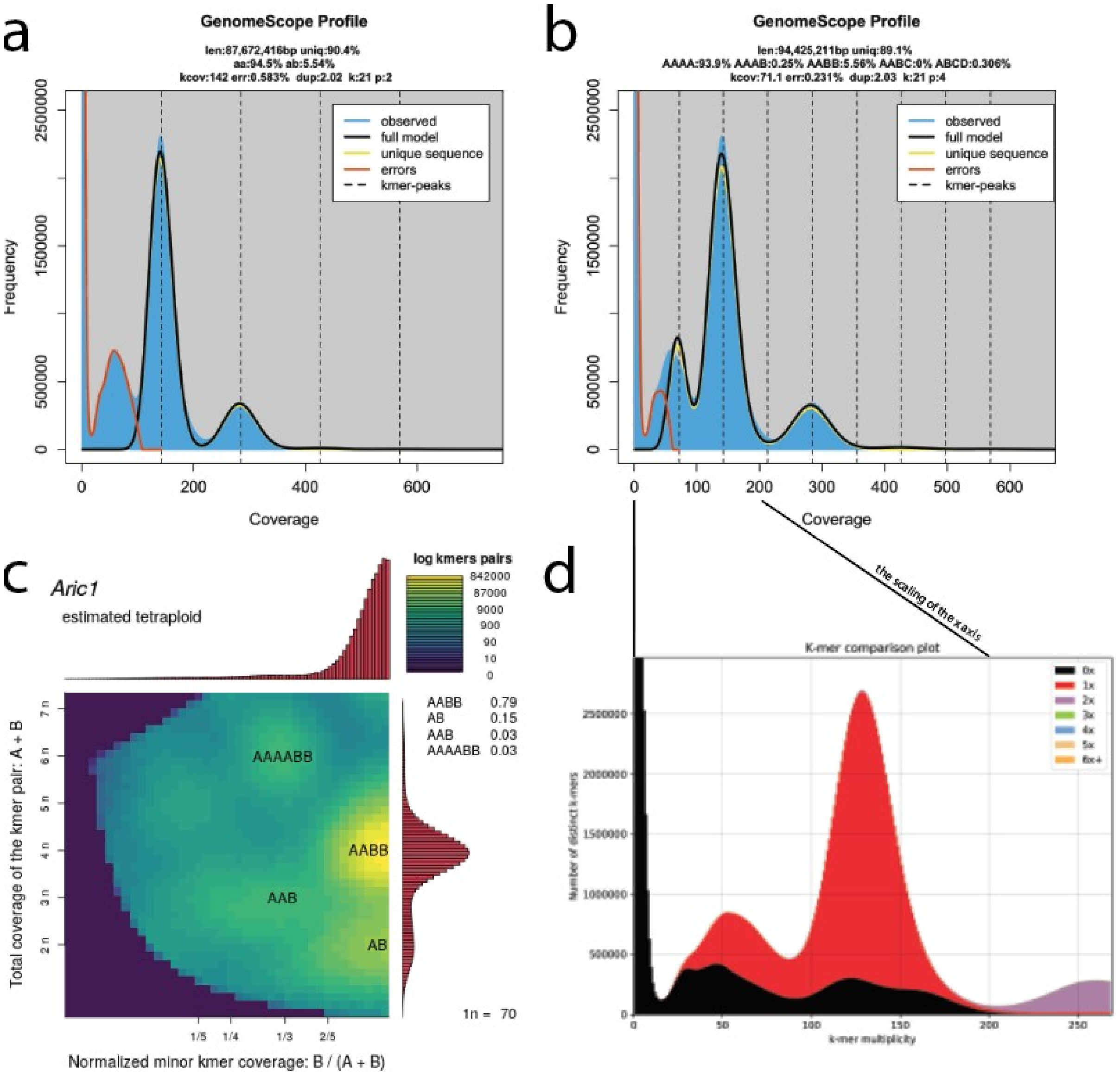


**Supplementary Figure 5: Genome profiling of *A. ricciae*. a** | Genome model used in our study. **b** | The best fit genome model supporting degenerate tetraploidy, which, assuming homoeologs have no shared kmers in the bdelloid rotifers, corresponds to degenerate octoploidy (for a more detailed interpretation see **Supplementary Material S2**) **c** | smudgeplot of *A. ricciae* indicating a 70x haploid coverage, and supporting the genome model shown on panel b. **d |** The comparison of the kmers in reads (i.e. the kmer spectra; x axis) and kmers in the decontaminated genome assembly (the coloration) suggest that kmers at least the 70x peak is part of the rotifer genome.

#### S3 Expected fraction of triallelic loci in triploid species

The observed biallelic and triallelic heterozygous loci can be indicative of the genome structure only when compared to an expectation. Assuming heterozygous alleles are randomly distributed across the genome we can generate a naive expectation of the fraction of triallelic loci in triploid species (in the absence of fitness effects linked to heterozygosity). This expectation is dependent on two variables: 1) how symmetric the pairwise divergences between the three genomic copies are, and 2) the total heterozygosity levels. In the main text (Figure 3) we report variation among triploid species in the frequency of bi- and triallelic loci. To verify that this variation is not solely generated by different total heterozygosity levels, we compared the observed proportion of triallelic loci among species while taking the bias generated by total heterozygosity levels into account. Consider a reference genome copy and two genome copies with divergences to the reference $d_1$ and $d_2$. The total heterozygosity (h) of the genome is $h = d_1 + d_2 - d_1d_2$ and can be further decomposed into biallelic heterozygosity ($d_1 + d_2 - 2d_1d_2$), and the triallelic heterozygosity ($d_1d_2$) as the overlap between the divergences $d_1$ and $d_2$. Note that the overlap is the highest in a genome with equidistant divergence of the genomic copies ($d_1 = d_2$). Given a heterozygosity *h*, the expected triallelic heterozygosity can be expressed as $d_{tri} = (1 - sqrt(1 - h))^2$.

This expectation does not correspond to the biological reality as genomes contain many regions with elevated or reduced heterozygosity but it allows us to compare genomes while correcting for the bias generated by different total heterozygosity levels. The frequencies of triallelic loci remain highly variable among the five triploid species even after correcting for this bias (**Supplementary Figure 6**), meaning the conclusions in the main text are supported even when we correct for total heterozygosity levels.

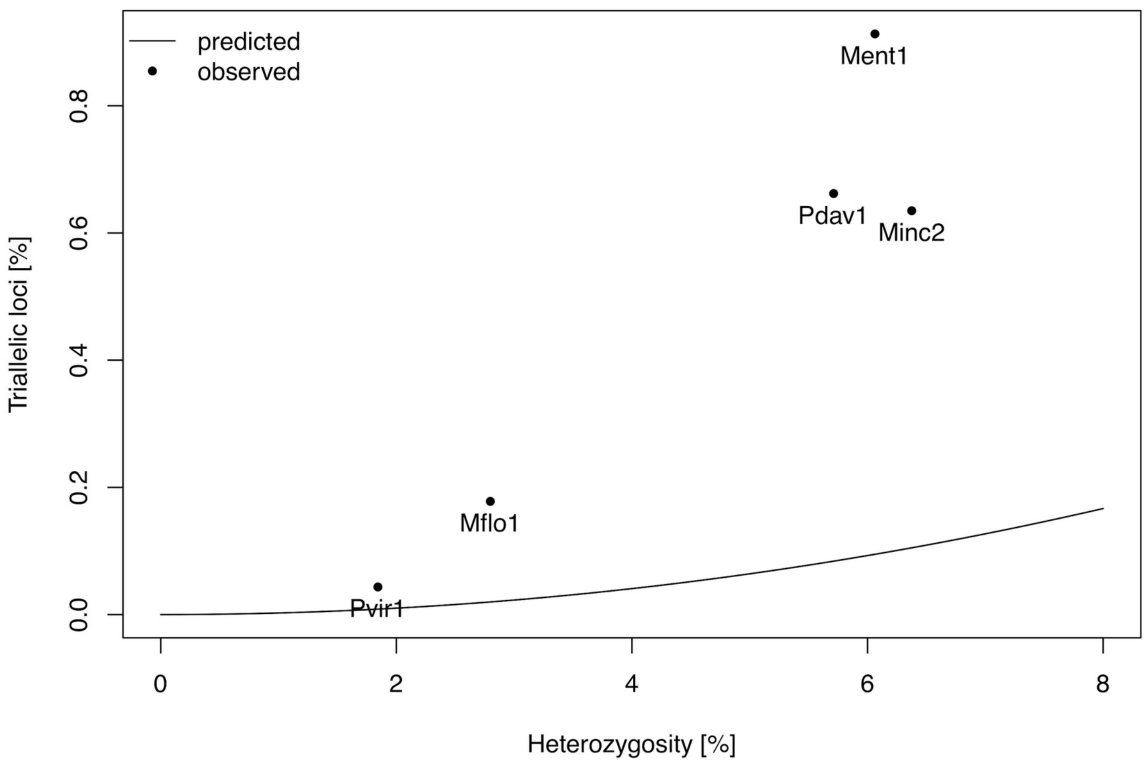


**Supplementary Figure 6: Fraction of triallelic loci according to the total heterozygosity.** We observe two distinct groups of triploid species. The crayfish (Pvir1) and *M. floridensis* (Mflo1) have a much smaller proportion of triallelic loci than the three other species, even when values are adjusted for a bias generated by total heterozygosity. The expectation was calculated as the overlap of heterozygosity in a genome with equidistant divergence (See **Supplementary Materials S3**).

#### S4 Suggestive hybrid origin of the marbled crayfish

Previously it has been suggested that the triploid parthenogenetic crayfish *Procambarus virginalis* is an autopolyploid lineage derived from diploid sexual *P. fallax* [117]. The main arguments for autopolyploidy (instead of allopolyploidy) are that *P. virginalis* and *P. fallax* are morphologically very similar and that *P. virginalis* does not carry any morphological trait of any other closely related crayfish. Our analysis revealed two nearly identical genome copies (**Figure 3**) supporting endoduplication (i.e., autopolyploidy) as a source of triploidy. However, it also revealed the presence of a highly diverged genome copy, suggesting hybridization between at least highly diverged strains or populations, if not species.

Specifically, the heterozygosity estimate for the parthenogenetic triploid *P. virginalis* is ~1.8% (**Supplementary Figure 7a**). Assuming endoduplication, this heterozygosity is generated by the third haplotype, diverged from the two identical copies. If so, we expect ~1.8% to also be the heterozygosity of sexual *P. fallax* individuals, the sexual sister species of *P. virginalis*. The heterozygosity of *P. fallax* is, however, much lower (~0.76%; **Supplementary Figure 7b**). This suggests that at least one of the haplotypes was acquired from a more diverged population via hybridization. However, to conclusively determine the origin of *P. virginalis*, we would need to better understand the population genetic diversity of sexual *P. fallax* and haplotype structures in *P. virginalis*.

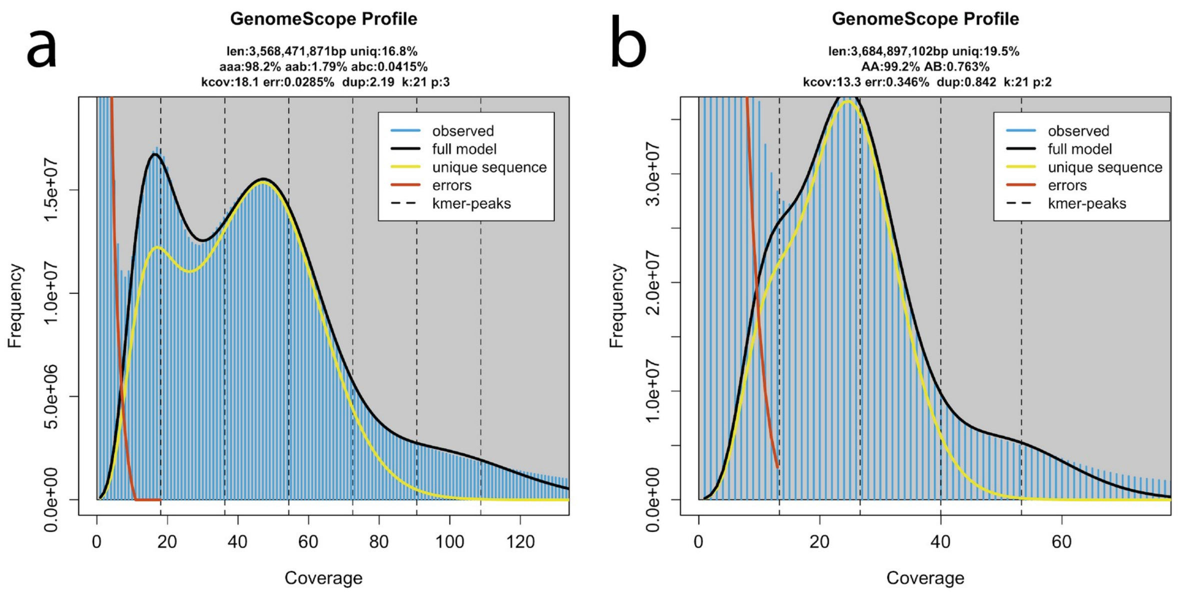


**Supplementary Figure 7: Genome profiling in crayfishes. a |** A triploid genome model in parthenogenetic *P. virginalis* estimates heterozygosity to 1.79%. **b |** A diploid genome model in its sexual sister species *P. fallax* estimates a similar genome size, but substantially lower heterozygosity (0.76%). However, the quality of fit is less conclusive as the error peak (red) and haploid peak (leftmost black) largely overlap.

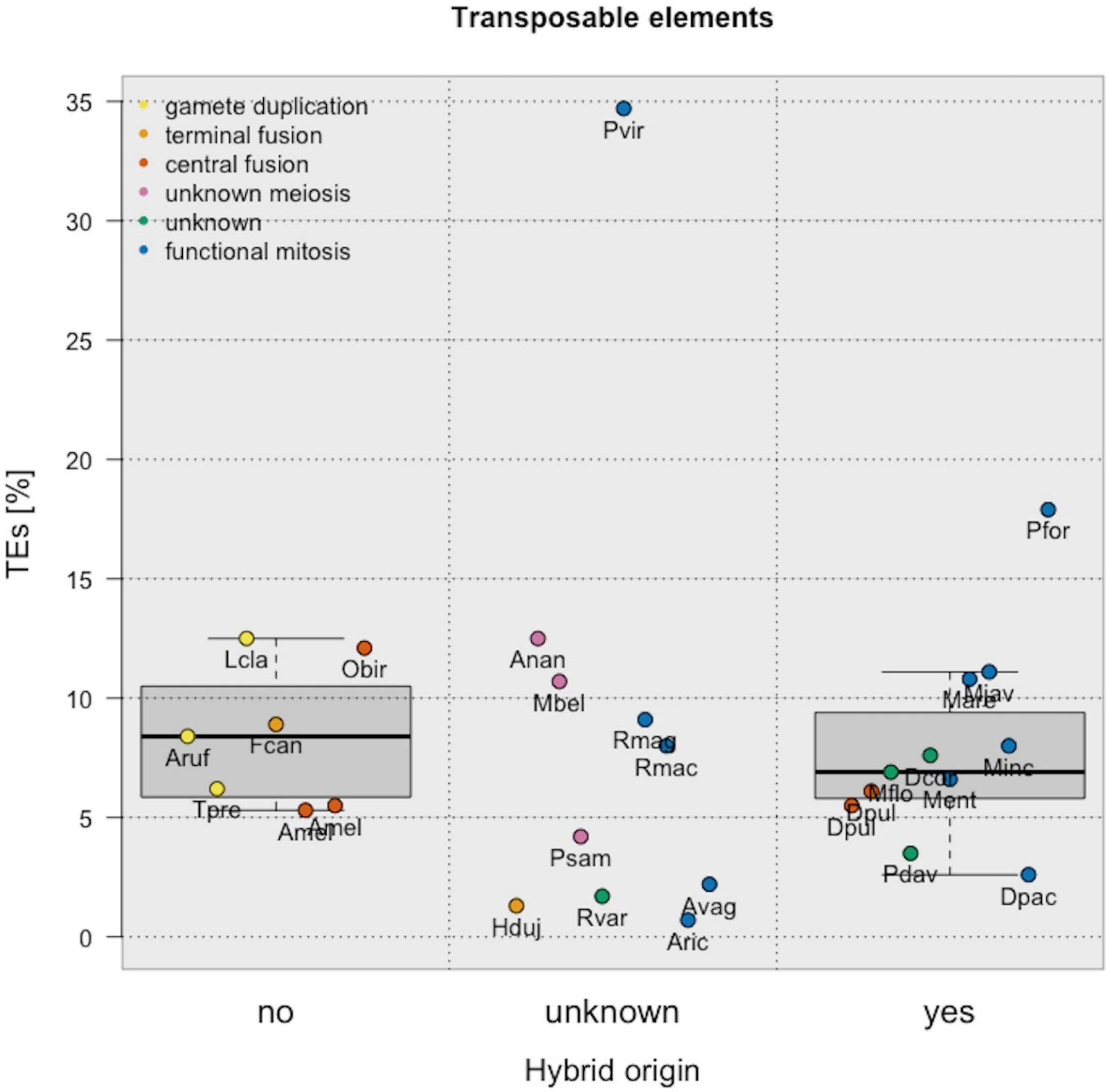


**Supplementary Figure 8: Transposable elements with respect to reproduction mode and hybrid origin.** Neither hybrid origin nor cellular mechanism of parthenogenesis are strong drivers of the TE content in parthenogenetic animals.

#### S5 Conserved gene content

We aimed to provide insights into gene duplications and losses by quantifying conserved single copy orthologs (BUSCO genes) [118]. BUSCO genes are defined as a set of genes that are present as a single copy in at least 90% of species inventoried in a curated database. All of the species used to build this database are sexual, and we initially hypothesised that both higher duplication rates and gene losses in parthenogenetic as compared to sexual species could be reflected in the percentages of missing and duplicated BUSCO genes in the analyzed parthenogenetic genomes. However, organisms that are highly heterozygous are prone to generating separate assembly of homologous haplotypes. In such split genome assemblies, BUSCO genes will falsely appear to be duplicated. To investigate whether split haplotype assemblies are of concern in the analyzed parthenogenetic genomes, we deduced the level of haplotype splitting in the assembled genomes by dividing the length of each assembly by the haploid genome size estimated from the read data with genomescope (higher frequencies of separate haplotype assemblies result in higher assembly length to haploid genome size ratios). We indeed found that BUSCO genes appear to be duplicated in genome assemblies consisting of split haplotypes, with the highest level of “artificial duplication” found in polyploid species of hybrid origin (**Supplementary Figure 9a**).

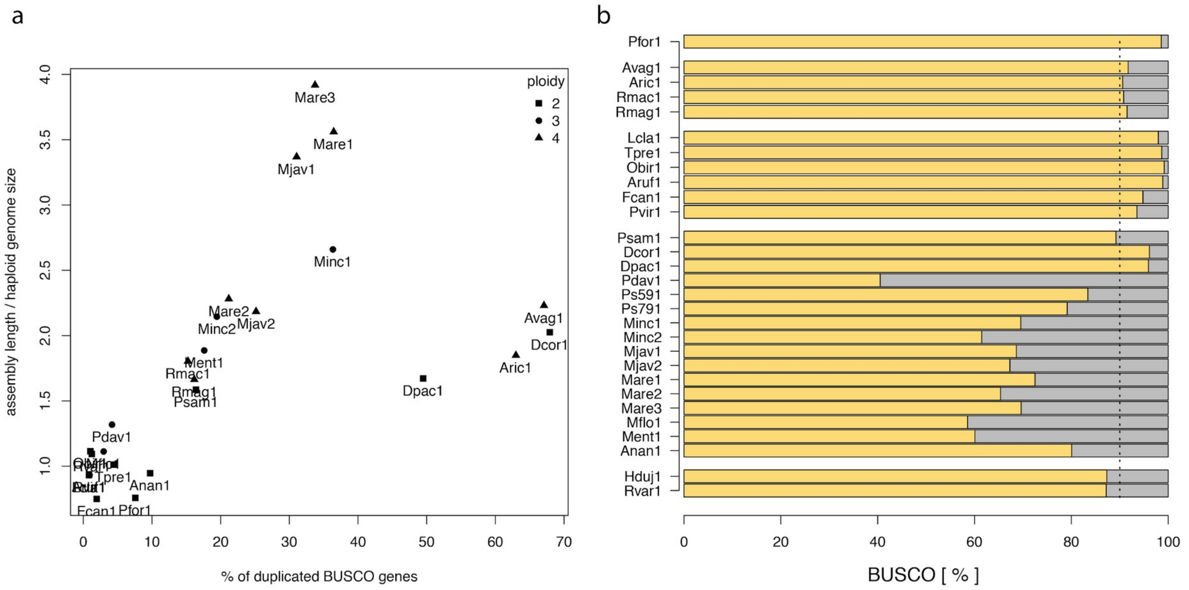


**Supp lementary Figure 9: Conserved single copy orthologs. a** | the fraction of duplicated BUSCO genes is correlated to the ratio of assembly length to haploid genome size. **b |** yellow bars show a proportion of BUSCO genes found in individual genomes. The dashed line indicates the expected level.

**Supplementary Table 1: Overview of analysed species.** This information was collected directly from the cited literature. References include information regarding cellular mode of reproduction, origin and/or the age of parthenogenesis.

https://github.com/KamilSJaron/genomic-features-of-parthenogenetic-animals/blob/master/LaTeX/SM_table_1_reproduction_modes.pdf

**Supplementary Table 2: Genomic features calculated from raw data.** We used unified methods to estimate basic genomic properties directly from sequencing reads. Ploidy was estimated using smudgeplot for all species but *A. vaga* (see section **Heterozygosity structure in polyploids** for details). Genome size, heterozygosity and repeats were estimated using GenomeScope. Repeats denote the fraction of the genome occurring in more than one copy. The classified repeats, TEs and other types of classified repeats, were estimated using DnaPipeTE.

https://github.com/KamilSJaron/genomic-features-of-parthenogenetic-animals/blob/master/tables/genome_table_infered_from_reads.tsv

**Supplementary Table 3: genome assemblies: size, number of scaffolds, N50, BUSCO, number of annotated genes.** Statistics were calculated from the published genome assemblies and genome annotations shared by authors. BUSCO genes were searched using the metazoan database for all the non-nematode species. Nematodes are notoriously known for the high turnover of genes and we therefore used nematode specific BUSCO genes. The number of annotated genes were calculated as the number of lines in the annotation with the tag “gene”.

https://github.com/KamilSJaron/genomic-features-of-parthenogenetic-animals/blob/master/tables/assembly_table.tsv

*The number of genes was extracted using the tag “mRNA” since the keyword “gene” was not in the annotation file of Diploscapter coronatus.

**Supplementary Table 4: Horizontal gene transfer analysis.**
HGT candidate genes identified from comparisons to UniRef90 (sheet 1) and UniProtKB/Swissprot (sheet 2) databases.

https://github.com/KamilSJaron/genomic-features-of-parthenogenetic-animals/blob/master/tables/JOH-2020-024.S4Table.HGT_sheet1_uniref.tsv

https://github.com/KamilSJaron/genomic-features-of-parthenogenetic-animals/blob/master/tables/JOH-2020-024.S4Table.HGT_sheet2_uniprot.tsv
